# LORA: a Polymorphic Multi-sample Long Read Assembly Pipeline

**DOI:** 10.64898/2026.01.06.697901

**Authors:** Dimitri Desvillechabrol, Rania Ouazahrou, Juliana da Fonseca Pipoli, Gerald Späth, Thomas Cokelaer

## Abstract

Genome assembly from long-read sequencing data has become a standard approach for resolving complex genomic regions and producing high-contiguity assemblies. However, the diversity of available assemblers, their varying performance across species, and the need for reproducible workflows present ongoing challenges.

We developed LORA, an easy-to-use and reproducible application for assembling genomes from long-read data. LORA integrates several well-established assemblers, including Canu, HiFiasm, Flye, and Unicycler, as well as more recent tools such as Necat and Pecat. It is implemented as a Snakemake pipeline to parallelize tasks and support seamless execution on both local machines and computing clusters.

LORA includes multiple quality assessment steps, interactive HTML reports for interpretation, BLAST-based taxonomic identification, and completeness evaluation. Together, these features provide users with a comprehensive view of assembly quality and potential problematic.

We illustrate the capabilities of LORA using datasets from bacterial genomes and unicellular eukaryotes, sequenced with both PacBio and Oxford Nanopore technologies, highlighting typical outcomes and common pitfalls encountered during long-read assemblies. LORA is distributed as part of the Sequana project, an open-source framework designed for reproducibility, maintainability, and straightforward deployment across computing environments.

## 1 Introduction

De novo genome assembly is a fundamental step in genomics [1], aiming to reconstruct complete genome sequences without the use of a reference. The advent of short-read sequencing platforms dramatically increased data throughput and enabled the development of new assembly algorithms [2, 3]. Nevertheless, assembling genomes from short reads remains challenging due to the sheer volume of data and the intrinsic structural complexity of many genomes. Although short-read technologies such as Illumina generate highly accurate sequences, their limited read lengths prevent the resolution of repetitive and low-complexity regions, often resulting in assemblies fragmented into many contigs.

The emergence of long-read sequencing technologies, including Oxford Nanopore Technologies (ONT) [4] and Pacific Biosciences (PacBio) [5], has transformed the field by enabling assemblies that span complex regions and frequently produce near-complete chromosomes in both prokaryotic and eukaryotic genomes [6, 7, 8, 9]. PacBio and ONT rely on different sequencing chemistries, leading to distinct performance characteristics. PacBio offers two main read types: Continuous Long Reads (CLR) and High Fidelity (HiFi) reads generated by circular consensus sequencing. CLR reads typically reach accuracies of 87-92% with N50 values of 30-60 kb, whereas HiFi reads exceed 99% accuracy with N50 values of 10-20 Kb. Typical throughput ranges from 50-100 Gb for CLR and 15-30 Gb for HiFi [10, 11]. ONT Sequencing, in contrast, produces reads that can range from 10-200 kb, with accuracy up to 98% depending on the chemistry and base calling. a MinION flowcell generally yields 0.5-20 Gb, while the high-throughput PromethION platform can exceed 100 Gb [10].

As long-read sequencing has matured, many assemblers have been developed to exploit these data, enabling the reconstruction of high-quality genomes, especially for haploid species [12, 13, 14]. Benchmarking efforts have compared their performance across diverse organisms and datasets [15, 16, 17]. Still, not all genomes assemble easily. Bacterial genomes, for instance, can be grouped into three classes based on their repeat content [18]: class I genomes contain only small repeats (e.g., rDNA operons, 5–7 kb) and can often be assembled into a few contigs. Class II genomes harbor numerous mid-sized repeats (e.g., insertion sequences *≤* 7 kb), requiring higher coverage and longer reads for closure. Class III genomes contain large repeats (*≥*7 kb), often phage-derived, such as tandem duplications and segmental repeats, which remain challenging even with long reads.

Diploid and polyploid genomes introduce additional complexity. Long reads improve haplotype reconstruction by spanning multiple heterozygous variants and enabling long-range phasing [19], a task largely intractable with shortread alone. However, long reads are not error-free: raw error rates of 5-15% in ONT and PacBio CLR data [20] can obscure true heterozygous sites, complicate phasing, and lead to haplotype switch errors. Distinguishing heterozygosity from sequencing noise remains challenging, although modern basecalling and variant-calling models, such as DeepVariant [21], have improved diploid variant detection. Hybrid strategies that combine long reads with accurate short reads [22] can mitigate these issues and have enabled telomere-to-telomere assemblies of large eukaryotic genomes, including humans [23], albeit at substantial computational cost.

Because genomic architectures vary widely, assembly strategies must be tailored to the biological context: prokaryotic vs. eukaryotic genomes, circular vs. linear structures, plasmid presence, or levels of heterozygosity. For example, Flye [13] offers a dedicated plasmid-assembly module, while Canu [12] assemblies of circular genomes often require post-processing circularization tools. In heterozygous eukaryotes, phasing tools such as Whatshap [19] are needed to separate allelic sequences. Coverage depth also plays a critical role: insufficient coverage leads to fragmented assemblies, while excessively deep coverage can complicate graph resolution.

Assessing assembly quality is equally important. BUSCO [24] evaluates completeness using conserved orthologs, while CheckM [25] provides lineage-specific completeness and contamination estimates for prokaryotic assemblies. Metrics such as N50, alignment rates, and coverage profiles further characterize contiguity and accuracy, guiding downstream annotation and comparative analyzes.

Although long-read assembly is now routine when sufficient coverage and read quality are available, outcomes remain highly dependent on input data characteristics and the choice of assembler. Comprehensive quality control, annotation, and reporting across multiple samples require substantial effort. Reproducibility is an additional challenge, as each step relies on many third-party tools, databases, and parameters.

To address these challenges, we introduce LORA (LOng Read Assembly), a reproducible, modular, and user-friendly Snakemake-based [26] pipeline developed within the Sequana project[27]. LORA provides an end-to-end solution for genome assembly from raw long-read data (FASTQ or BAM), supporting both PacBio and ONT platforms. It integrates multiple assemblers, optional polishing with short reads, and thorough quality assessment using BUSCO, CheckM, Quast, and sequana-coverage [28], with results consolidated in rich HTML reports. Rather than proposing a new assembler or performing an exhaustive benchmarking study, LORA offers a standardized, versatile, and reproducible workflow that simplifies tool selection, parameterization, and reporting across diverse genomic contexts.

In the following sections, we describe the LORA workflow (Materials and Methods), evaluate its performance and usability on representative datasets (Results), and discuss its applicability and limitations across diverse genomic contexts (Discussion).

## 2 Materials and Methods

### 2.1 Genome sequencing data and genome assembly data

To demonstrate the versatility of the LORA pipeline, we examined a variety of datasets covering both ONT and PacBio technologies, spanning different genomic complexities.

#### 2.1.1 PacBio datasets

PacBio, sequencing has evolved rapidly, producing different data types with distinct accuracy profiles. In this study, we first used FastQ files derived from the Continuous Long Read (CLR) data. More recent sequencers generate Consensus Circular Sequence (CCS) reads, including *HiFi* reads, obtained by performing multiple passes of the same molecule and applying a stringent quality filter (at least 10 passes, predicted accuracy of 99%, denoted *mp* and *rq* respectively). While final CCS/HiFi reads are typically distributed as FASTQ files, the underlying data are stored in PacBio-specific BAM files. These BAM files differ from standard alignment BAMs: they encode raw subreads, per-base qualities, polymerase metadata, and the information required to reconstruct CCS or HiFi sequences.

#### 2.1.2 ONT datasets

For ONT, we considered several cases, including multi-sample datasets and data produced with different chemistries and basecalling models, such as the recent R10.4 model. In this study, we used data in FastQ format obtained after basecalling.

#### 2.1.3 Genomic diversity

Regardless of the sequencing technology used, we analysed genomes with varying repeat architecture and levels of biological complexity. Bacterial genomes can be classified into three repeat-defined classes according to [18]. Using 100,000 bacterial genomes, which were downloaded from the NCBI and are available via Zenodo entry odo.11519700, we updated this classification. Genomes with sizes below 1 Mbp were removed, leaving 42,000 genomes for analysis. Repeats were identified with the Repeats class from the Sequana library [27]. Most genomes fall into class I and class III (96%), and these classes form the core of our long-read datasets. To broaden the applicability of LORA, we additionally considered a small eukaryotic genome, whose chromosomes can similarly be classified as class I and class III.

### 2.2 Datasets

#### 2.2.1 *Veillonella Parvula* – CLR PacBio data

As a representative of Class I genomes, we selected *Veillonella parvula* SKV38, which contains only short repeats (below 1.5 kb; see Annex). Sequencing reads (ERR3958992) were obtained from the Sequence Read Archive (SRA) in FastQ format and published in [6]. The genome was sequenced on a PacBio Sequel II platform and assembled into a single contig of 2,146,482 bp (NCBI accession NZ LR778174).

The dataset includes N=338,310 reads with an average length of L=9 kb and an N50 of 13 kb. The depth of coverage (*N ×L/*genome size) is therefore approximately equal to 1, 500 *X*. To minimize misassemblies, the authors randomly subsampled 100,000 subreads, corresponding to 600 *X* coverage. Assembly was performed with Canu v1.8, circularized using Circlator [29], and polished with Illumina data. The final assembly (RefSeq GCF 902810435.1) reports 98.36% completeness and 1.66% contamination according to CheckM [25]. The original BAM files are also available on Zenodo, enabling the reconstruction of CCS reads (DOI 10.5281/zenodo.13306683).

#### 2.2.2 *Streptococcus Gallolyticus subsp. Macedonicus (SGM)* – HiFi PacBio data

As an example of a Class III genome, we analyzed *Streptococcus gallolyticus subsp. macedonicus* (SGM), which is characterized by large repeats of 6 kb and 10 kb located in close proximity (see Annex). Sequencing reads (SRR24332397) were obtained from the SRA and published in [30] as part of the PRJNA94176 project. The final assembly (GCF 029866925.1) comprises a 2,210,410 bp chromosome and a 12,729 bp plasmid.

The dataset associated with SRR24332397 (from PRJNA940176) contains approximately 2.5 million subreads in FASTQ format, generated on a PacBio Sequel II platform. In the original publication, the authors performed assembly using CCS reads generated with a minimum accuracy of 99% and at least three passes. However, these CCS reads are not available in the SRA, and the corresponding BAM file required to reproduce them was not publicly deposited.

To enable the generation of HiFi reads, we obtained permission from the authors to redistribute the original subreads BAM file. This file is now available on Zenodo (DOI: 10.5281/zenodo.17207091). Using this BAM file, we reconstructed HiFi CCS reads with the desired parameters (mp *≥*10 passes, rq *≥* 99%). For convenience, both the HiFi BAM file and its corresponding FASTQ version are also provided in the Zenodo entry.

#### 2.2.3 *Bacteroides Fragilis* – multi samples Nanopore data

To demonstrate LORA’s capability to handle multiple samples, we reanalyzed data from Sydenham et al. [31], where several assemblers were benchmarked on Bacteroides fragilis isolates sequenced with PacBio, Nanopore, and Illumina technologies.

The reference genome (ASM2598v1, NCBI accession NC 003228.3) is categorized as Class I, with six 5 kb repeats and only a few 1-3kb repeats. However, the study isolates contained much large repeats (*≥* 7kb, up to 20kb), spanning both Class I and III categories (See Figure 2).

**Figure 1:**
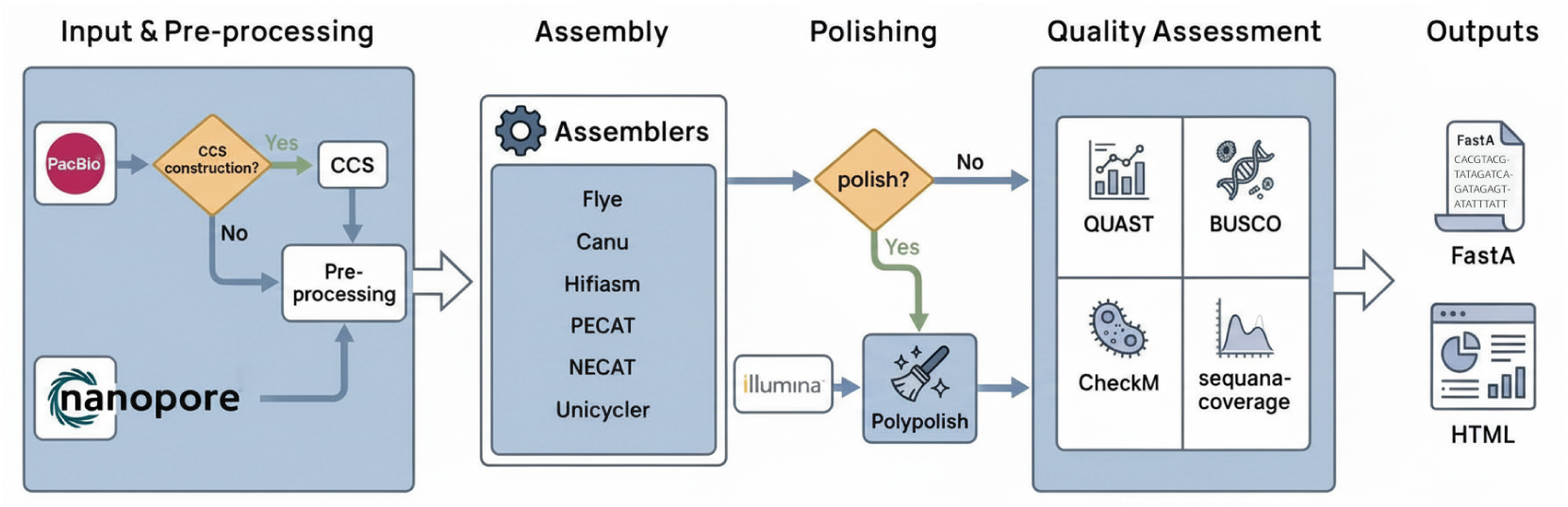
Overview of the LORA pipeline. Input and pre-processing accepts PacBio (BAM/FastQ) and ONT (FastQ) data. Can automatically generate high-quality CCS/HiFi reads from PacBio BAM files. Asssembly employs one of six integrated assemblers chosen by the user. Polishing is optional and can refine the assembly accuracy using short-read Illumina data. Quality assessment is made of a comprehensive suite of tools (BUSCO, Quast, CheckM, sequanacoverage) to evalutte contiguity, completeness, and contamination. The outputs delivers assembled contigs (FastA) and rich interactive HTML reports for easy interpretation.

**Figure 2:**
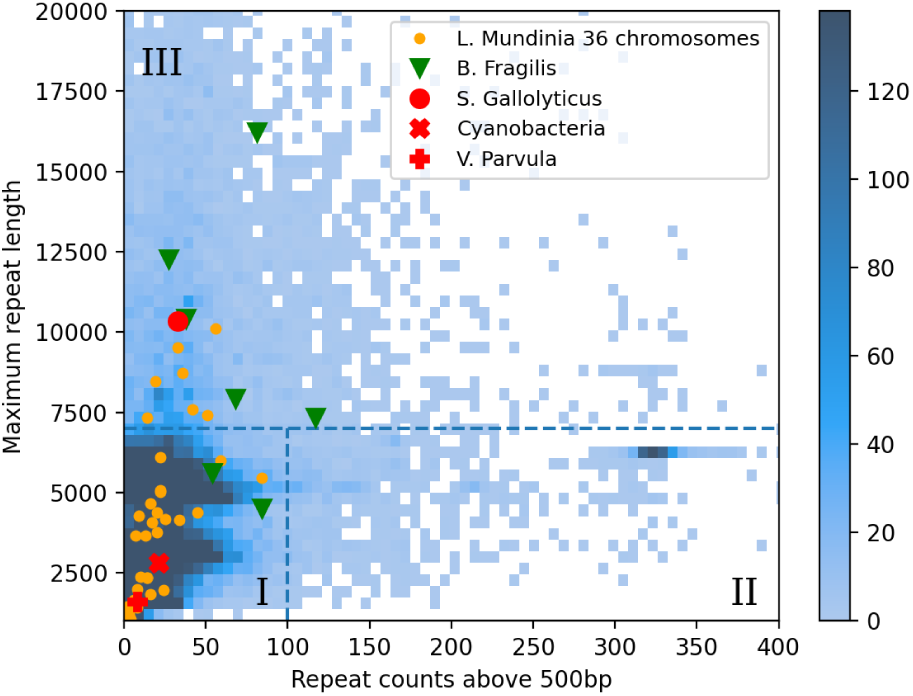
Repeat count versus maximum repeat length. This density plot includes 44,600 bacterial genomes of at least 1 Mb. For clarity, values above 20 kb for the repeat count are clipped, retaining 97% of the original dataset. Using a threshold of 7 kb for the maximum repeat length and a threshold of 100 for the number of repeats longer than 500 bp, three classes of genomes are defined. Classification and plot inspired by [18].

Nanopore (R9.4.1) FastQ files are available via Zenodo entry zenodo.2677927). Additional data, including raw signal files (fast5 format), PacBio reads, and Illumina data are accessible under BioProject accession numbers PRJNA525024, PRJNA244942, PRJNA244943, PRJNA244944, PRJNA253771, PRJNA254401, and PRJNA254455.

The reference genome assembly mentioned above is available under NCBI accession number GCF000025985 and reports a completeness value of 99% and total length of 5,205,140 bp. A plasmid, pBF9342, is also available (NC 006873.1; 36,560 bp).

#### 2.2.4 *Cyanobacteria* – contamination Nanopore data

To showcase LORA’s integrated quality assessment tools, we analyzed an unpublished dataset of *Cyanobacteria* that included contamination from two distinct strains. Sequencing was performed on a GridION device using R10.4 chemistry and high-accuracy base-calling model.

The final assembly yielded two chromosomes of 4.41 Mb and 4.13 Mb. Repeat structure suggests a Class I classification. BLAST identified one chromosome as *Limnospira* and the other as *Geminocystis*. Data is available via Zenodo (DOI 10.5281/zenodo.17229238)

#### 2.2.5 *Leishmania* – eukaryotes Nanopore

Finally, to evaluate LORA’s applicability to eukaryotic genomes, we analyzed a *Leishmania* dataset from Almutairi et al. [32]. *Leishmania* genomes are diploid (often aneuploid), contain 34-36 chromosomes, and have a total genome size of approximately 33 Mb, making them compact but structurally complex eukaryotic genomes.

We focused on the *Leishmania (Mundinia)* sp. Ghana strain, for which ONT and Illumina datasets are available (NCBI project PRJNA691536). Long read data is provided by Nanopore (R9.4.1) FastQ files. The genome was assembled using a Snakemake-based workflow [32], yielding 125 contigs with an N50 of 961,565 bp (NCBI assembly GCA 017918215.1).

### 2.3 LORA pipeline

The analysis presented in this study was performed using LORA (version 1.0.0), a modular and extensible assembly pipeline. LORA relies on sequana_pipetools (version 1.0.3; sequana_pipetools), which provides the command-line interface for pipeline execution (See Section Discussion for details). The LORA source code and documentation are available within the Sequana project organization at github.com/sequana/lora. An overview of the LORA workflow is shown in Figure 3. The main features of LORA are detailed in Section Results.

**Figure 3:**
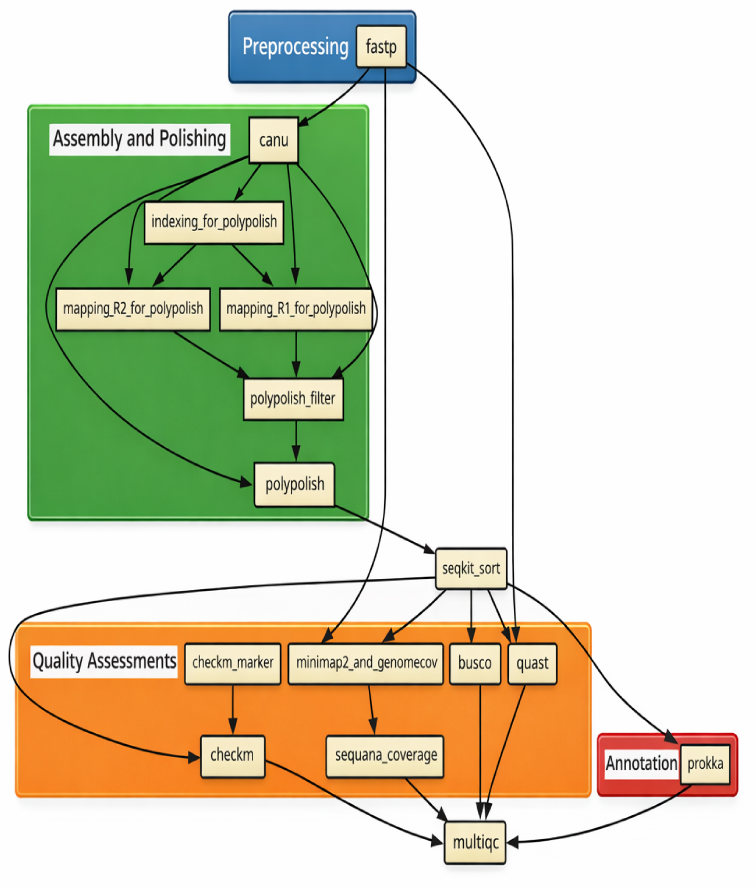
Example of LORA workflow. This workflow begins with input FASTQ data (blue box), followed by assembly with Canu and polishing using Illumina reads (green box). The resulting contigs are then subjected to quality assessments (orange box) and annotation (red box). Final results consist of HTML reports and contigs in FastA format.

## 3 Results

### 3.1 Pipeline overview

LORA is a modular and reproducible Snakemake-based pipeline designed for long read genome assembly. It is part of the Sequana project [27], which provides a comprehensive suite of next-generation sequencing (NGS) pipelines covering genomics, transcriptomics, and both short- and long-read technologies.

At its core, LORA leverages Snakemake [26], a workflow management system that combines the readability of Python with a flexible domain-specific language. Snakemake ensures transparent rule definition, automatic handling of dependencies, and efficient use of computational resources through built-in parallelization and portability across different computing environments, from laptops to high-performance clusters.

Beyond Snakemake, LORA benefits from the sequana_pipetools library), which provides a unified interface across all Sequana pipelines. This framework standardizes configuration handling, command-line options, and report generation. As a result, users familiar with one Sequana pipeline can readily adopt others, and developers can maintain consistency across the ecosystem. The user interface is deliberately kept simple: each pipeline can be launched with a single command, while advanced parameters can be fine-tuned via a configuration file compatible with Snakemake.

A simplified version of LORA is shown in Figure 1. The full workflow, including task dependencies, is illustrated in Figure 3, which presents a rule graph with specific dependencies. Tasks (or “rules” in Snakemake terminology) at the same level are essentially independent and can be executed concurrently.

Moreover, the pipeline has a polymorphic design, adapting its structure based on the user’s input data and chosen options. For example, Figure 3 includes optional tasks such as polishing with Illumina data or performing quality assessments.

In the following sections, we provide a detailed description of LORA’s components including supported assemblers, quality assessment modules, and reporting features.

### 3.2 Reporting

Once LORA completes its analysis, all results are organized in a clear directory tree. Each sample has its own subdirectory, which in turn contains folders for each task (e.g., preprocessing, assembly, polishing). To make it easy for users to explore the outputs, LORA generates a set of HTML reports. The first report is produced with MultiQC [33], providing multi-sample summaries for tools such as fastp, BUSCO, sequana coverage, and others. LORA also provides its own entry point (summary.html), which links to the MultiQC report, the pipeline configuration file, its dependencies, an overview of the workflow, and a summary table for each sample, including key metrics such as the number of contigs, and links to Quast reports. The summary page also links to a more complete report that includes links to individual contig coverage plots. When BLAST analysis is enabled, the report also includes information on the best BLAST hits for each contig. If enabled, it also contains the BUSCO and Checkm metrics.

### 3.3 Input Data

LORA accepts long-read sequencing data in standard FASTQ format, supporting both PacBio and ONT datasets. For ONT, only base-called FASTQ files are supported; raw signal formats such as pod5 or fast5 are not handled directly. PacBio data may be provided in FastQ format or BAM files. Although BAM files can be converted to FASTQ with third-party tools (e.g., using BioConvert [34]), LORA natively supports them by converting them into FastQ files.

PacBio sequencing generates subreads, i.e. single passes over a DNA molecule, which typically exhibit an error rate of 10–15%. To reduce these errors, PacBio can perform multiple passes over the same DNA molecule and generate a consensus sequence (CCS) known as HiFi reads. HiFi data usually require at least ten passes and can achieve an accuracy of 99%. When users provide original subreads in BAM format, LORA can automatically generate HiFi or other CCS reads. This step is efficiently incorporated into the workflow by splitting the input BAM files, computing consensus sequences, and merging the resulting data back into a single BAM file.

### 3.4 Preprocessing

Once the input data are available as FastQ file, provided directly by the user or through the building of the CCS data (PacBio only), LORA processes the data using FastP [35] allowing for quality control and filtering. By default, reads shorter than 1,000 bp are filtered out, but users can customize this threshold and apply additional FastP options, such as quality trimming.

### 3.5 Assemblers included

The pipeline integrates several assemblers that have different strengths depending on the nature and quality of the input data.

#### 3.5.1 Canu

The Canu assembler [12] is known for its ability to manage highly erroneous reads and produce high-quality assemblies. Canu implements an overlap–layout–consensus (OLC) approach first described in Rodger Staden in 1979. Canu includes dedicated modules for read correction, trimming, and assembly. A genome size estimate can be provided by the user, although this parameter is not directly enforced during contig construction. Circularisation is not included and should be performed with other tools such as Circlator [29] to close circular chromosomes or plasmids. It does not explicitly model diploidy; heterozygous regions are typically collapsed into a consensus sequence.

#### 3.5.2 Flye

Flye assembler [13] assembles genomes using a repeat graph, which accommodates high sequencing error rates and captures the genome’s repeat structure. This approach enables the recovery of long contigs even from noisy long-read data. Flye can optionally circularize replicons internally, making explicit post-processing unnecessary in many cases. For diploid genomes, Flye generally produces a collapsed mosaic haplotype rather than fully phased sequences, though the original assembly graph can be retained for downstream haplotype analysis or inspection.

#### 3.5.3 Hifiasm

LORA also includes the Hifiasm [36] assembler that is optimised for PacBio HiFi reads and produces high-quality, haplotype-resolved assemblies. It begins with an all-versus-all overlap of reads, followed by several rounds of error correction, then constructs a string graph where bubbles represent heterozygous allelles. Depending on user options and parental data, Hifiasm can create a primary/alternate contigs or fully phased haplotypes. Circularisation is not an explicit feature, as Hifiasm is primarily targeted at linear chromosomes and phased eukaryotic assemblies.

#### 3.5.4 NECAT

NECAT [37] is a de novo assembler designed primarily for ONT, though it is compatible with PacBio CLR data. It adopts an Overlap–Layout–Consensus (OLC) strategy with a progressive error-correction scheme: low-error reads are corrected first, followed by high-error reads. This adaptive procedure accelerates correction while preserving accuracy. During assembly, NECAT constructs contigs from the corrected reads and subsequently bridges them with raw reads to improve continuity. An adaptive selection mechanism is used both to identify high-quality supporting reads for each template during correction and to retain the most reliable overlaps in the layout stage. NECAT can assemble bacterial, fungal, and moderately sized eukaryotic genomes on standard hardware. It does not include an internal circularisation step, and heterozygous regions are generally collapsed rather than phased. Compared with “correct-then-assemble” tools such as Canu, NECAT is reported to be 2.5–258 × faster while producing assemblies of similar quality; relative to “assemble-then-correct” assemblers such as Flye, NECAT achieves fewer misassemblies on complex genomes [37].

#### 3.5.5 PECAT

Another assembler included in LORA is PECAT [38] (Phased Error Correction And Assembly Tool). It is also specifically designed for long, noisy reads (e.g. Nanopore or PacBio CLR) to reconstruct diploid genomes with haplotype-aware accuracy. It follows a correct-then-assemble approach: a haplotype-aware error correction step that preserves heterozygous alleles followed by two rounds of string-graph-based assembly. The second step uses SNP-calling components to filter inconsistent overlaps and rescontruct haplotype-specific contigs, either in “primary/alternate” or in “dual assembly” format. PECAT achieves very high continuity (e.g. large N50) and compares favorably to other diploid-capable assemblers. It is efficient in CPU time and memory, relative to more classical correct-then-assemble pipelines, and demonstrates good performance even with noisy reads, especially as coverage and read quality increase. PECAT does not include built-in circularisation, as its focus is on accurate reconstruction of phased contigs in diploid or polyploid contexts.

#### 3.5.6 Unicycler

Unicycler [14] is a de novo genome assembler developed primarily for bacterial genomes, supporting short-read, long-read, and hybrid assembly modes. It integrates the short-read assembler SPAdes [3] with a long-read bridging algorithm, enabling the resolution of repetitive regions and the closure of circular chromosomes or plasmids when long reads are available. Although Unicycler can assemble Illumina data alone, long reads alone, or a combination of both, LORA currently supports only the long-read mode. In this mode, Unicycler employs minimap and miniasm to construct an initial assembly, which is uncorrected and therefore retains an error rate similar to that of the input reads. The bundled version of miniasm is modified to better assemble circular replicons into circular string graphs. Unicycler also incorporates polishing steps (e.g., Racon) to refine consensus accuracy and improve genome continuity. Owing to its graph-based architecture, Unicycler is particularly well suited to small, low-complexity genomes but may be less efficient for large or highly heterozygous eukaryotic datasets [14].

### 3.6 Post processing

#### 3.6.1 Circularisation

After the assembly step, bacterial genomes and plasmids often require an additional circularization step to represent their natural topology. Assemblers may output a linear contig even when the underlying molecule is circular, leaving duplicated ends or small overlaps. To address this, LORA includes a circularization module based on Circlator[29] that scans contig termini for significant overlap, trims redundant regions, and reorders the sequence so that the origin of replication or another user-defined position appears at the start. This procedure ensures that circular chromosomes and plasmids are accurately represented as single, gap-free sequences, facilitating downstream analyses such as annotation and comparative genomics. Circlator is currently not maintained (sanger-pathogens/circlator). A fork was created within the Sequana project to fix several bugs. This version has been containerised and is distributed through the Damona project as a reproducible container.

#### 3.6.2 Scaffolding

Assemblers generally output a set of contigs, representing contiguous stretches of sequence but lacking information about their relative order and orientation. When a reliable reference genome is available, these contigs can then be aligned, generating a scaffold. This alignment organizes the contigs into the correct order and orientation, while introducing placeholder gaps (denoted by N bases) for regions that remain unresolved. LORA integrates the RagTag suite [39] to perform this scaffolding step efficiently, mapping contigs to the reference and generating a scaffolded assembly that is ready for downstream analyses.

### 3.7 Quality assessments

Quality assessment is a critical step in evaluating the accuracy, completeness, and reliability of a genome assembly. Several key metrics are commonly employed to assess assembly quality.

#### 3.7.1 N50

The N50 metric is a widely used metric to assess genome assembly contiguity. It is defined as the length *N* such that 50% of the total assembly length is contained in contigs (or scaffolds) of length greater than or equal to *N* . To calculate it, contigs are first ordered from longest to shortest, and their lengths are cumulatively summed until half of the total assembly size is reached. The length of the contig at this threshold is reported as the N50 value. While N50 is most informative for assemblies with many contigs, it can be less meaningful for assemblies generated from long-read data with only a few contigs. In LORA, Quast [40] is used to compute the N50.

#### 3.7.2 Alignment rates

This measures the proportion of reads from the original long-read sequencing data that successfully map to the assembled contigs. High alignment rates indicate that a large fraction of the sequencing data has been incorporated correctly into the assembly. In LORA, Quast [40] is used to compute the alignment rate.

#### 3.7.3 Completeness

Another useful metric is completeness. It reflects the proportion of a genome successfully recovered in the assembly. It is typically estimated by identifying conserved, single-copy marker genes that are expected to be present in all members of a particular taxonomic group. LORA integrates BUSCO (Benchmarking Universal Single-Copy Orthologs) [24], which assesses assembly completeness by comparing the genome to a set of highly conserved, single-copy genes. A high BUSCO completeness score indicates that most essential genes have been successfully captured.

LORA also includes CheckM [25], which provides a completeness estimate for bacterial genomes and offers finer resolution across sub-taxonomic groups.

#### 3.7.4 Contamination

This refers to the presence of sequences from other organisms within the assembled genome. This can occur when metagenomic sequences from different species are incorrectly combined during assembly. Both BUSCO and CheckM provide contamination estimates. BUSCO identifies duplicated marker genes that may indicate contamination from closely related strains. CheckM distinguishes between two types of contamination: (i) fragments originating from multiple strains of the same species, and (ii) fragments from more distantly related taxa (heterogeneity). High heterogeneity can suggest the presence of sequences from different strains or subspecies.

#### 3.7.5 Genome Coverage

Depth of coverage along the genome is useful to evaluate how much of the target genome is represented in the assembly. Higher coverage generally correlates with higher completeness. In LORA, genome coverage is computed using sequana coverage [28], which maps reads to contigs, estimates coverage trends, and provides visualizations for inspecting contigs, detecting copy number variations (CNVs), and identifying coverage drops.

Collectively, these quality assessment metrics offer a comprehensive view of assembly performance, helping researchers gauge the reliability and utility of the genome assembly. A high-quality assembly should exhibit high contiguity, completeness, accuracy, and low contamination, making it a valuable resource for downstream genomics and bioinformatics analyses.

### 3.8 Annotation option

#### 3.8.1 Prokka

LORA is dedicated to assembly. However, in the context of bacterial genomes, we can use Prokka [41] to annotate the final contigs. Currently, no annotation is planned for eukaryotes, which usually requires specific domain knowledge.

#### 3.8.2 Blast

LORA optionally supports the annotation of assembled contigs through BLAST searches against user-supplied reference databases. Because the size and scope of BLAST databases vary, depending on the intended application, LORA does not provide or distribute reference datasets. Instead, users are responsible for preparing and maintaining a valid local BLAST database prior to execution. When such a database is available and properly configured, LORA performs BLAST queries for each contig and incorporates the most relevant hits into its HTML report, thereby facilitating the inspection of putative annotations and improving the interpretability of the assembly.

## 4 LORA Pipeline Use Cases Across Diverse Genomes

In this section, we illustrate the versatility and flexible design of the LORA pipeline. Figure 3 shows a representative workflow; however, the actual execution may vary depending on the user’s setup, input data, selected mode (bacterial or eukaryotic), or choice of assembler. The datasets used in the examples below are described in section 2 and summarized in Table 1. They span different sequencing technologies and varying data quality over time.

**Table 1:**
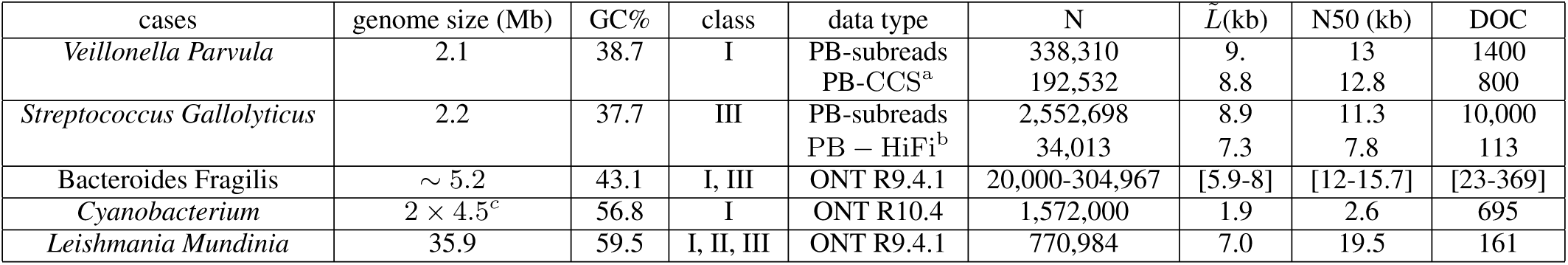
Data Summary. First column lists the species used on each test case. Column 2 reports the approximate genome length, followed by the expected GC content and the genome’s complexity class. For the *Leishmania* dataset, the complexity class was determined for each of the 36 chromosomes, which fall into three distinct categories. Additional information is provided for each dataset, including the sequencing data type, from either PacBio or Nanopore (for PacBio, datasets may consist of raw reads, CCS reads, or HiFi reads). The table then lists the number of reads in the raw or processed data, the mean read length (*L̃*), the N50, and the expected depth of coverage (*N×L/*genome size). Notes: *^a^* Default parameters of the LORA pipeline were used (rq *>* 0.7, mp *≥* 0). *^b^* HiFi reads were generated with rq *>* 0.99 and mp *≥* 10. *^c^* Due to contamination, two cyanobacterial genomes of approximately 4.5 Mb each were present.

### 4.1 *Veillonella Parvula* – CLR PacBio data

This case evaluates LORA’s ability to handle noisy CLR reads and compare assembler sensitivity to read quality. The original dataset, composed of subread files in BAM format, can be used directly by the pipeline, which automatically converts them to FASTQ files. Alternatively, the BAM files may be used to build circular consensus sequences (CCS) for downstream assembly.

The genome under study (*Veillonella Parvula*) is classified as class I and should not be difficult to assemble. Nevertheless, the dataset considered was sequenced on a PacBio Sequel II platform with an older chemistry version (v1.2) without multiplexing. As a result, the data yield was large (Table 1), but each molecule was sequenced only a few times. Using LORA, we first attempted to generate HiFi reads from the subreads; however, only a limited number of subreads met the HiFi quality criteria. In the original study [6], the authors used a subsample of 100,000 raw subreads (coverage 400*X*) without constructing CCS reads. Following this strategy, we performed two experiments: (i) assembling a random subsample of 100,000 subreads, and (ii) generating CCS reads by disabling filtering on the number of passes (setting the minimum to 0) and requiring an accuracy threshold of 0.7, yielding 175,000 CCS reads.

For both datasets, we ran all 6 assemblers available in LORA (Section Assemblers included) with and without the circularization option, resulting in 32 individual pipeline executions. Default parameters were used, and the `--mode bacteria` option was enabled to run BUSCO (bacterial lineage) and CheckM, using *Veillonella parvula* as the reference species. Assembly and completeness results are shown in Figures 4 and 6.

**Figure 4:**
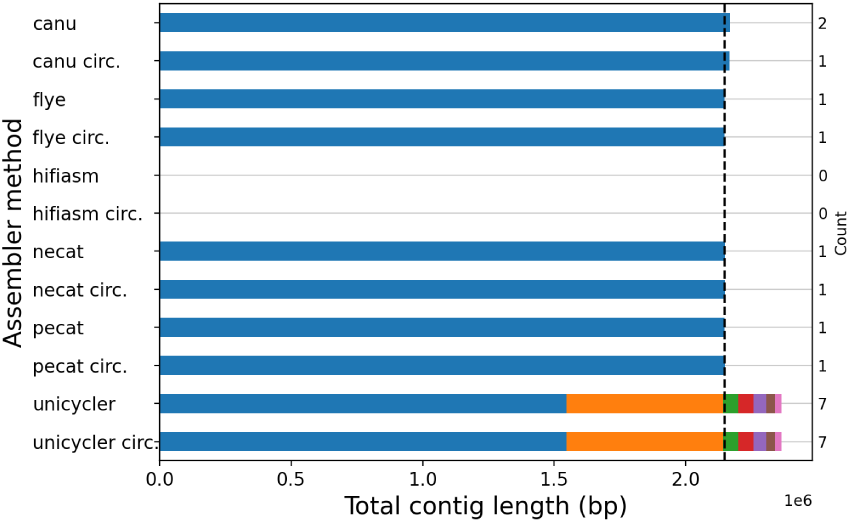
Assembly results for *Veillonella Parvula* using PacBio raw reads. Each horizontal bar represents the total size of the assembled contigs (X-axis), with the corresponding number of contigs shown on the right Y-axis. The assembler used, along with whether circularization was applied, is indicated on the left Y-axis. The vertical dashed line indicates the expected total size. Overall, most assemblers successfully completed the assembly, with the exception of Hifiasm, which failed to finish the assembly task. Unicycler produced seven contigs. Colors are used solely for visual distinction and carry no specific meaning.

**Figure 5:**
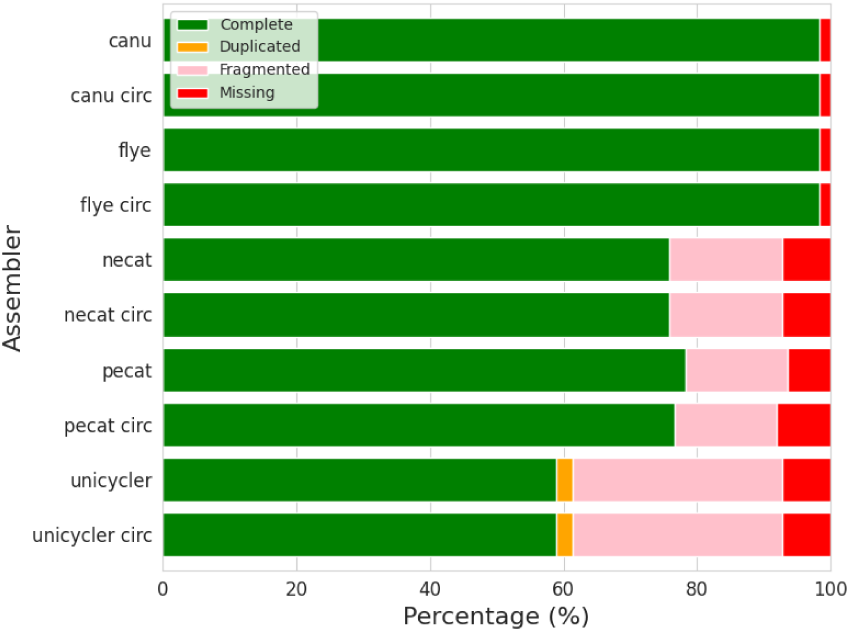
BUSCO scores obtained for the *Veillonella* case (subreads). The the Y-axis, the assemblers used for this analysis and on the X-axis, the Busco Score. While Canu and Flye achieved high completeness (bacteria lineage), others exhibits significant proportion of fragmented or missed genes.

**Figure 6:**
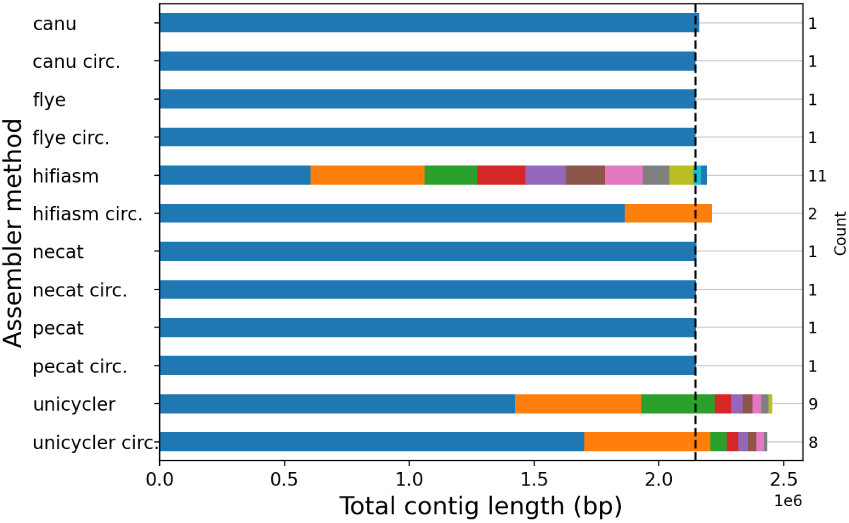
Assembly results for *Veillonella Parvula* using PacBio CCS reads (see text for details). Each horizontal bar represents the total size of the assembled contigs (X-axis), with the corresponding number of contigs shown on the right Y-axis. The assembler used, along with whether circularization was applied, is indicatd on the left Y-axis. The vertical dashed line indicates the expected total size. Overall, most assemblers successfully completed the assembly, with the exception of Hifiasm and Unicycler that generated about 10 contigs. Colors are used for distinctions of the contigs

**Figure 7:**
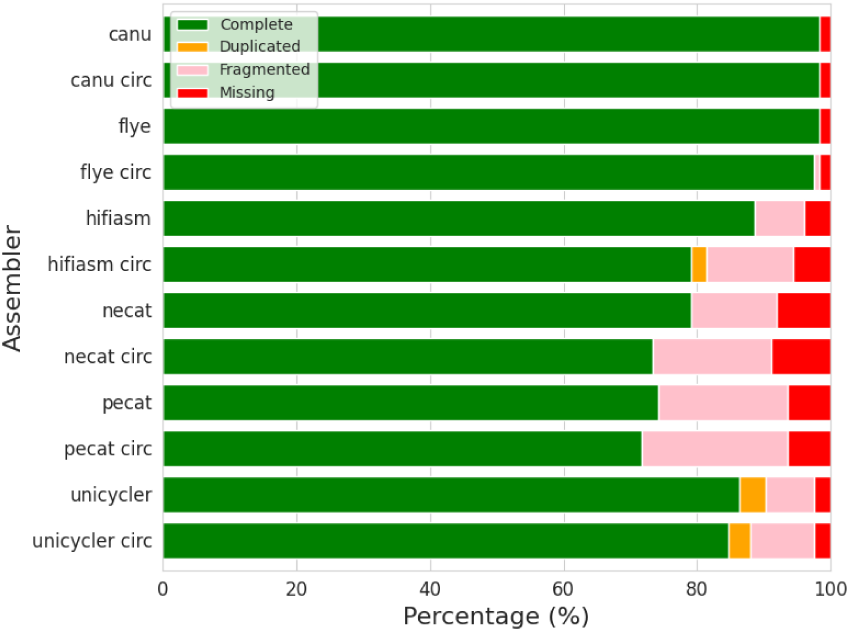
BUSCO scores obtained for the *Veillonella* case (CCS). On the Y-axis, the assemblers used for this analysis and on the X-axis, the Busco Score. While Canu and Flye achieved high completeness (bacteria lineage), others exhibits significant proportion of fragmented or missed genes.

Using the subsampled subreads, most assemblers achieved a good assembly, as shown in Figure 4. Canu produced a single contig when the circularization option was enabled; without circularization, the same contig was obtained, but an additional small contig was reported (not supported by coverage). CheckM completeness reached 98.5%. Flye produced one complete contig with or without circularization. Flye correctly identified the origin of replication even without explicit circularization, indicating robust internal handling of circular genomes (CheckM: 98.4%). Hifiasm failed on this dataset, likely due to the high error rate in the raw reads. Necat generated a single chromosome, but quality control exposed the low quality of the assembly (BUSCO: 9 missing and 21 fragmented genes; CheckM: 93%), with several local coverage drops. Pecat yielded a single chromosome, with similar issues (8 missing genes and 19 fragmented BUSCO genes). Circularization slightly improved CheckM (*≈* 95%) but not the BUSCO scores. Unicycler performed poorly, producing seven contigs, a CheckM score of 97.5%, and only 76 complete BUSCO genes.

When using CCS reads, coverage increased, and read quality improved (*≈*175,000 CCS reads). Canu produced results similar to those obtained from raw reads, consistent with its internal correction step (CheckM: 98.4%). Flye again achieved an excellent assembly (CheckM: 98.4%, BUSCO: 112/124), with no benefit from circularisation. Hifiasm finished its tasks but remained unsuitable, although enabling circularisation reduced the number of contigs from seven to two but introduced substantial coverage drops (CheckM: 95%). Necat assembled a complete chromosome but retained low CheckM completeness (93%) and fragmented BUSCO genes. Pecat also reconstructed the chromosome with the correct length but yielded CheckM completeness of only 94.5% and irregular coverage. Unicycler produced seven contigs with a total size exceeding the expected genome length by *≈* 20%.

In summary, Flye and Canu consistently produced high-quality assemblies, with Flye generally yielding the most accurate and stable results, independent of whether circularisation was applied. In contrast, Hifiasm was sensitive to read quality, and Unicycler tended to over-fragment the assembly. Necat and Pecat were able to generate single-chromosome assemblies, but with reduced completeness and irregular coverage. These results demonstrate that LORA enables users to systematically benchmark assemblers and parameter choices for optimal genome reconstruction.

### 4.2 *SGM* – HiFi PacBio data

Here we test LORA’s performance on repeat-rich, high-quality HiFi datasets. We considered a bacterium classified as class III due to a 10ḱb repeat and several 6 kb repeats, notably near the origin of replication (Figure 22). Starting from the raw data, we set the pipeline to build the PacBio HiFi CCS using the `--pacbio-build-ccs` option. This is different from [30] where CCS reads were generated with *≥* 3 passes and an accuracy above 99%.

After running the pipeline for each assembler, both with and without circularization, all assemblers completed the assembly. A summary of the number of contigs obtained for each assembler is shown in Figure 8, and all BUSCO scores are provided in Figure 9.

**Figure 8:**
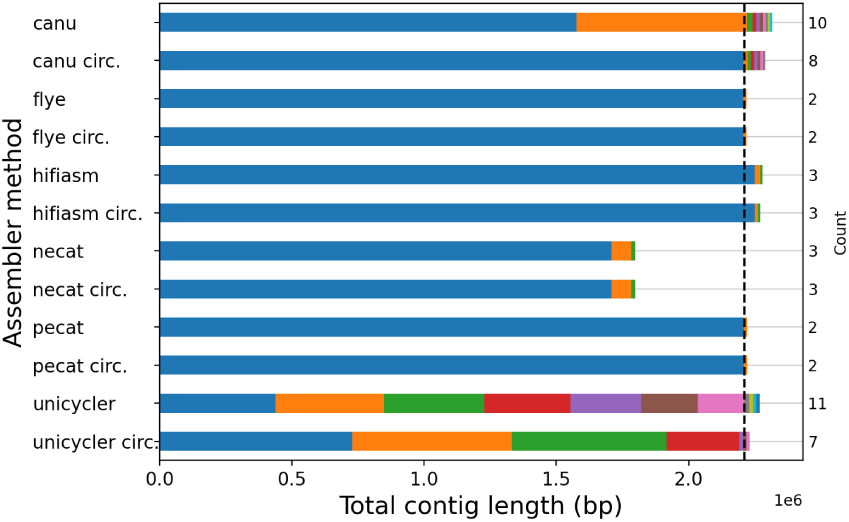
*SGM* Assembly results using HiFi CCS reads with 110X coverage for *SGM*. Each bar correspond to the number of contigs obtained (Y-axis, right) and it’s size (x-axis), depending on the assembler used (Y-axis, left). The expected genome consists of one full chromosome and one plasmid. All assemblies successfully recovered the plasmid. Canu generated 10 contigs and, Unicycler 11 contigs, each one represented by its respective color on the graph.

**Figure 9:**
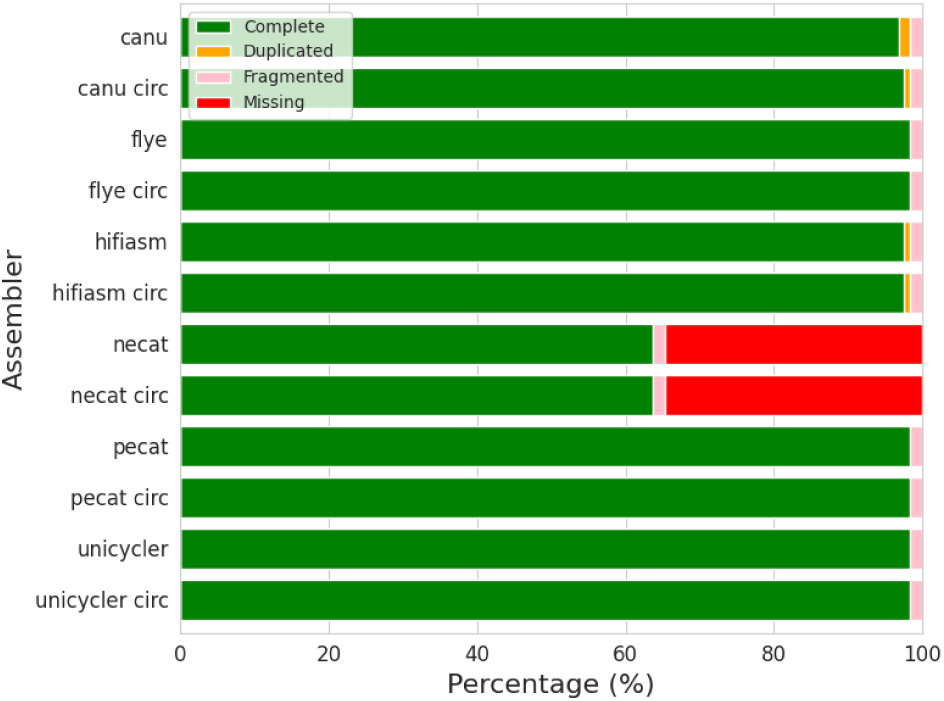
BUSCO scores obtained for the SGM case. On the Y-axis, the assemblers used for this analysis and on the X-axis, the Busco Score. Except for necat, all assemblers achieved a high completeness on the bacteria lineage.

**Figure 10:**
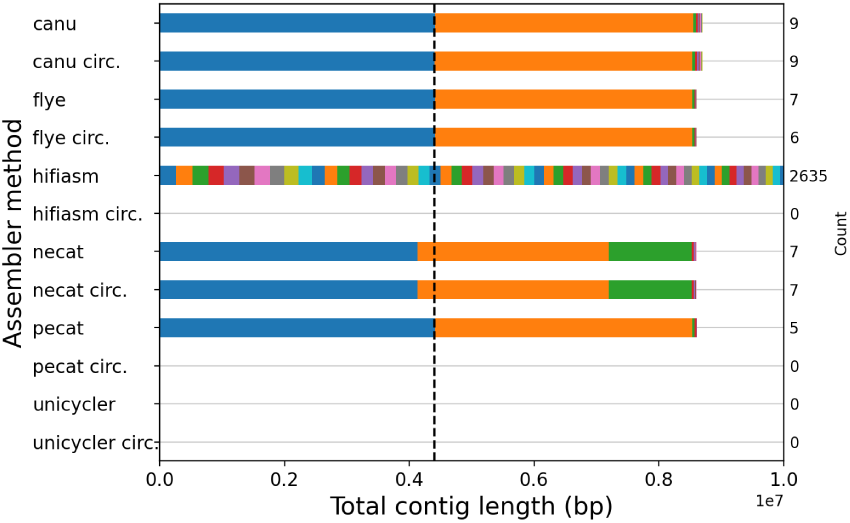
*Cyanobacteria* Assembly results with 366X coverage for *cynobacterium* test case. Each bar correspond to the number of contigs obtained (Y-axis, right) and it’s size (x-axis), depending on the assembler used (Y-axis, left). The expected outcome is 2 full chromosome of 2 different cyanobacteria and several plasmids. Most assemblers found the two species and plasmids, represented by different colors on the graph. Hifiasm circ., Necat circ., Unicycler failed the task and did not finish. Hifiasm generated more than 2000 contigs. Metrics such as CheckM and BUSCO scores are reported as percentages or counts, showing genome completeness, contamination, and errors. * means plasid found but incorrect length

**Figure 11:**
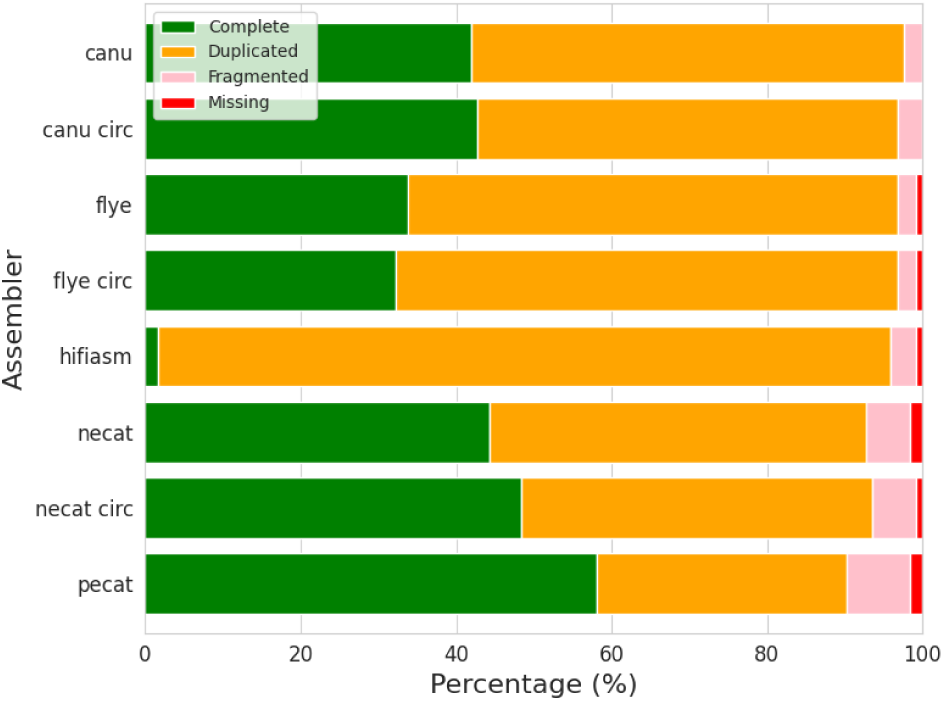
BUSCO scores obtained for the *Cyanobacteria* case. On the Y-axis, the assemblers used for this analysis and on the X-axis, the Busco Score. Due to the contamination with two closely related bacteria, the duplicated rate reach 50% in most cases.

Combined with a circularization step, Canu produced an almost perfect assembly comprising a circular chromosome and plasmid. Canu also assembled six small artefactual contigs, that can be easily discarded. The plasmid showed only minor local drops in coverage when the dataset was mapped to the assembly. If we omit the circularization step, the chromosome is split into two contigs and the plasmid is incompletely resolved.

Flye consistently returned two high-quality contigs representing the chromosome and plasmid. The circularization step mainly adjusted the origin of replication without altering continuity.

With circularization, Hifiasm yielded a complete chromosome and plasmid, plus a single unsupported contig that can be ignored. Without the circularization step, the plasmid assembly was inaccurate (excess length).

Assemblies produced by NECAT, irrespective of the circularization step, were incomplete: the largest contig covered about 80% of the chromosome, accompanied by a 70 kb fragment and a misassembled plasmid.

PECAT delivered results comparable to Flye. The circularization step slightly reduced overall quality, introducing minor coverage dips.

With and without the circularization step, Unicycler generated fragmented, yet complete assemblies (7–11 contigs) and the plasmid was recovered with high precision.

Overall, Flye and PECAT provided the most straightforward and accurate reconstructions, while Canu (with circularization) also performed well after filtering small contigs. Hifiasm results were also quite good regarding contiguity and completeness because of the high quality of the data set. NECAT and Unicycler were clearly less suitable for this genome, and circularization had tool-specific effects, generally improving assembly orientation but occasionally affecting coverage uniformity.

### 4.3 Cyanobacterium – Nanopore R10 with contamination

This case demonstrates LORA’s capacity to disentangle mixed genomes and identify contamination. We use a recent high-yield Oxford Nanopore dataset. Although the sequencing experiment targeted a single cyanobacterium, contamination was present, resulting in two species (estimated genome sizes 4.1 Mb and 4.4 Mb) together with two plasmids. This dataset is particularly interesting because it enables an assessment of how assemblers available in LORA behave in complex, mixed samples.

Canu, without circularisation, successfully reconstructed the two bacterial chromosomes with uniform coverage. CheckM reported 99.2% completeness but also a contamination level of 90%, while BUSCO revealed that over half of the genes were duplicated, reflecting the presence of both species. Plasmids were detected but not fully assembled. Enabling circularisation in Canu yielded similar results, with improvements on fully closing the circular chromosomes.

Flye produced highly contiguous assemblies in both modes. Without circularisation, it recovered the two chromosomes with 99.5% completeness, alongside four additional contigs: a correctly assembled 35 kb plasmid, a small 12 kb plasmid with proper 400X coverage, a 14 kb fragment with abnormally high coverage ( 8000×), and another lowquality sequence. Applying Flye’s circularisation step did not change these results, indicating that internal handling of circular elements is already effective.

Necat (with or without circularisation) assembled one of the genomes as a single 4.1 Mb contig, while the second was split into two pieces. Completeness was slightly lower than Flye (*≈* 99%), and coverage plots showed localized drops. Both plasmids were retrieved, along with the same spurious high-coverage contig observed with Flye.

Pecat reconstructed the two chromosomes and plasmids when run without circularisation, achieving 98.5% completeness, though some coverage dips remained. However, pecat with circularisation failed to complete. Unicycler was unable to finish (memory usage exceeded 64 GB), and hifiasm generated an excessive number of contigs; its circularisation step did not converge, likely because of the fragmented assembly graph.

Overall, Flye offered the most balanced outcome for this contaminated Nanopore dataset, providing near-complete assemblies for both cyanobacteria and the plasmids, with minimal artefacts. Canu also performed well, while Necat and Pecat produced usable results but with slightly reduced completeness. Assemblers such as Unicycler and Hifiasm were not suited for this challenging mixed sample.

### 4.4 Isolates of Bacteroides fragilis

This case illustrates LORA’s multi-sample management and integrated polishing features. We use a Bacteroides dataset consisting of seven samples sequenced with both Oxford Nanopore Technologies (ONT), PacBio CLR, and Illumina short reads. This dataset was used to demonstrate (i) the ability of LORA to process multiple samples in a single run, and (ii) the integration of polishing steps based on short-read data. To handle multi-samples, users simply need to indicate where to find the input files (`--input-directory`) and a pattern (default uses the wildcard *.fastq.gz). For polishing, LORA incorporates the Polypolish tool [42]. Similarly to the long read data, users need to indicate where to find the FastQ files (`--polypolish-input-directory`). LORA handles single-end and paired-end data sets.

Assemblies were first generated from ONT data using Flye. Across the seven samples, assemblies ranged from 3 to 11 contigs with a total length of 5.4 Mb *±* 0.15 Mb. Coverage varied from 20X to 200X, with lower coverage samples producing a higher number of contigs. Some assemblies included circularized replicons, such as sample S01 (a 5.5 Mb chromosome and a 55 kb plasmid) and nBF02 (three circular contigs corresponding to the chromosome and two plasmids). Overall, the assemblies were structurally sound. However, quality metrics revealed limitations: CheckM completeness was 81% and BUSCO scores (Figure 12) indicated fragmentation (41%) and missing genes (15%), consistent across all seven samples.

**Figure 12:**
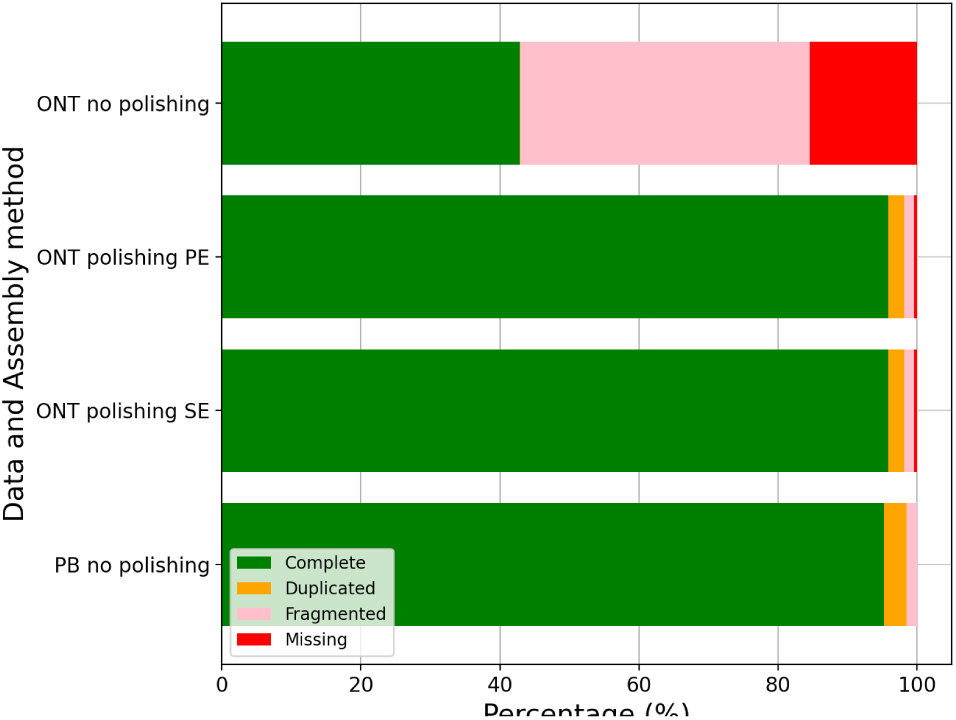
BUSCO scores obtained for for the *Bacteroides fragilis* case with and without polishing

Assemblies were then repeated with PacBio CLR data at 100× coverage. In this case, final assemblies were more fragmented with 10–40 contigs, with a total length of 5.5 Mb *±* 0.2 Mb. However, the consensus accuracy was higher. Here, CheckM completeness reached 99% and BUSCO completeness was 98.5%, a striking contrast to the 43% observed with ONT-only assemblies.

Finally, to demonstrate LORA’s integrated polishing functionality, we combined ONT assemblies with Illumina pairedend reads. Two polishing modes were tested: one using paired reads and one using only the first mate. In both cases, the overall contiguity of assemblies remained unchanged, but polishing improved gene completeness: BUSCO completeness increased to 98.2% and CheckM completeness to 99%, demonstrating the effectiveness of short-read polishing in correcting long-read assemblies.

Within LORA, one HTML output is made with MultiQC[33]. BUSCO plugin is integrated within lORA. An instance is shown in Figure 13.

**Figure 13:**
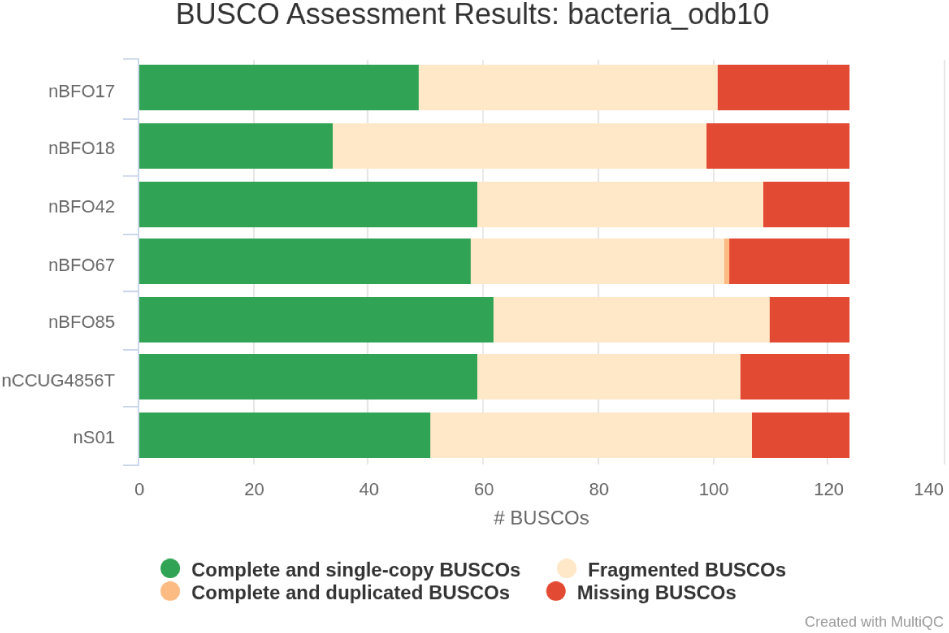
BUSCO completeness scores for the Bacteroides assemblies (ONT case). This MultiQC plot, automatically generated within the LORA pipeline, summarizes the proportion of complete, fragmented, duplicated, and missing genes across the seven samples.

**Figure 14:**
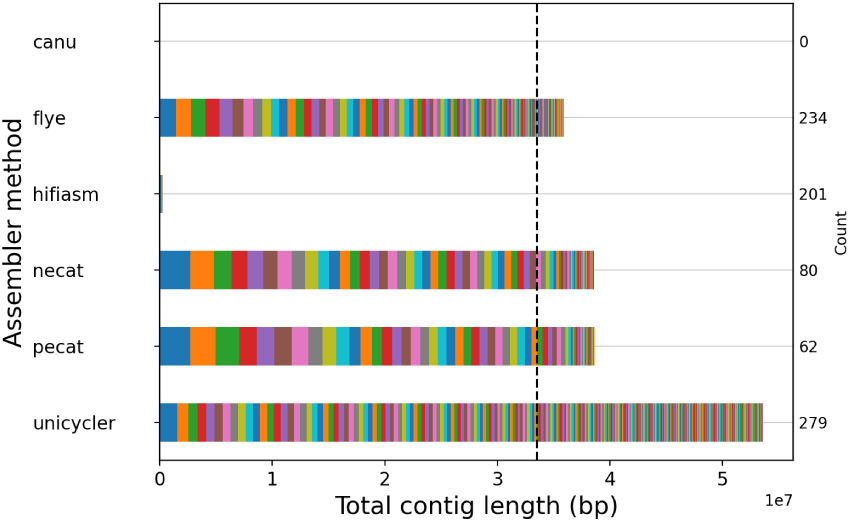
Assembly results for *Leishmania* dataset. Each horizontal bar (divided into colored segments) correspond to the contigs obtained by different assemblers (Y-axis, left). The number of contigs is provided (Y-axis, right) and their total length indicated on the X-axis. The expected outcome is 36 full chromosomes, 1 maxi circle (35 kb) and several mini circles (1 kb)for a total of about 35 Mb. Canu correction was stopped after 3 days of computation. Total sum of contigs produced by hifiasm is below 250,000 bases.

**Figure 15:**
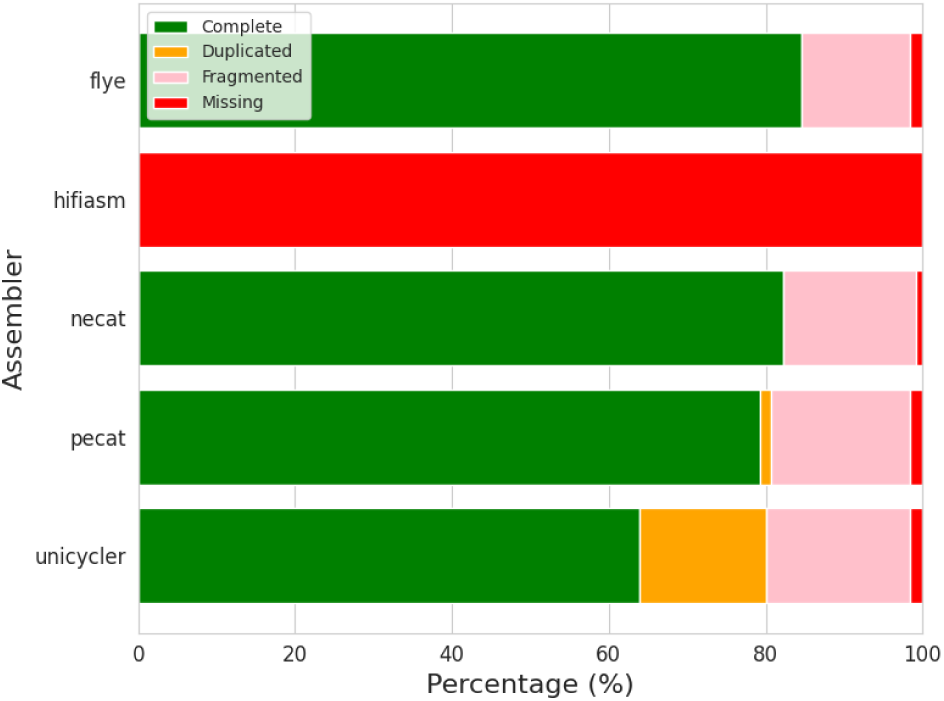
BUSCO scores obtained for the *Leishmania* case. The Y-axis shows the assemblers used in this analysis, and the X-axis indicates the BUSCO score (percentage). Flye, NECAT, and PECAT exhibit similar results, with a large proportion of complete genes and some fragmented ones. Unicycler shows an excess of duplicated genes, while hifiasm misses most genes due to assembly failure.

### 4.5 Applying LORA on a small eukaryote genome from *Leishmania*

Finally, we assess LORA’s applicability to small diploid eukaryotic genomes. We considered the genome of *Leishmania Mundinia*, a kinetoplastid parasite with a complex genome. This organism is typically diploid, although not all chromosomes are strictly diploid, and its genome comprises 36 chromosomes totaling approximately 33 Mb. We used low-coverage Oxford Nanopore long reads as input as described in Materials and Methods.

Although LORA ships with assemblers designed primarily for bacterial genomes (e.g., Unicycler), it also integrates tools capable of managing more complex assemblies, including diploid eukaryotes. The aim of this case is not to benchmark assemblers exhaustively but to demonstrate that LORA supports a variety of tools suitable for eukaryotic genomes.

Among the tested assemblers, Canu failed to produce an assembly (more than 48h computation) and Hifiasm failed to produce meaningful contigs (200 contigs of length below 3kb). Flye generated 234 contigs spanning 36 Mb, with a largest contig of 1.5 Mb. In contrast, NECAT achieved better contiguity, producing 80 contigs, including a 2.7 Mb contig corresponding to the complete chromosome 36. PECAT performed similarly well, yielding 62 contigs for a total size of 38.6 Mb, with the same chromosome recovered as a single contig.

LORA also provides an optional scaffolding step using a known reference. When applied to NECAT and PECAT assemblies, this feature reconstructed all 36 chromosomes with only 16 and 14 gaps, respectively. Both NECAT and PECAT assemblies reached an N50 of 800 kb, compared to 500 kb for Flye. However, NECAT and PECAT showed slightly more fragmented BUSCO genes (+3 relative to Flye) and an increase of 3 Mb in assembled sequence, suggesting a trade-off between contiguity and assembly accuracy.

As expected, Unicycler, which is optimized for bacterial genomes, struggled with this dataset: it produced 279 contigs totaling 53 Mb, likely reflecting partial phasing of the diploid genome. BUSCO analysis confirmed 20% duplicated genes, consistent with overrepresentation of heterozygous regions.

Overall, this case study highlights that LORA’s ecosystem of assemblers extends beyond bacteria. With appropriate tool selection and optional scaffolding, users can assemble small eukaryotic genomes such as *Leishmania* with satisfactory results, even under limited sequencing coverage.

## 5 Discussion

### 5.1 Philosophy

LORA was designed to provide a unified, reproducible, and transparent framework for genome assembly using longread sequencing data. Rather than introducing a new assembler, LORA’s main objective is to orchestrate existing best-in-class tools, allowing users to easily compare their performance under consistent and reproducible conditions. The case studies presented in this work demonstrate that assembly outcomes strongly depend on both the input data type and genome complexity. By integrating multiple assemblers, preprocessing steps, and quality assessment modules into a single automated pipeline, LORA enables users to identify the most suitable strategy for their specific datasets.

### 5.2 Reproducibility and FAIR principles

LORA was designed to comply with the principles of reproducible research and the FAIR guidelines (Findable, Accessible, Interoperable, and Reusable) [43]. The workflow is fully version-controlled and documented, ensuring transparency and traceability of all analyses. To guarantee reproducibility across computing environments, LORA uses containerized software dependencies. All non-Python tools, such as minimap2, BUSCO, CheckM, and the included assemblers, are distributed through the Damona project https://damona.reathedocs.io as Apptainer (formerly Singularity) containers [44]. These containers encapsulate all necessary binaries and libraries, eliminating system-level variability. When LORA is executed, the required containers are automatically downloaded and managed by the pipeline, requiring no manual installation or configuration. This approach ensures consistent execution across local machines, HPC clusters, and cloud environments, while simplifying deployment for users. By combining transparent configuration, versioned workflows, and containerized dependencies, LORA offers a robust and reproducible framework for long-read genome assembly, fully aligned with FAIR data and software principles.

### 5.3 Benchmarking philosophy

LORA is not intended to benchmark assemblers exhaustively but rather to facilitate orchestrated evaluation under standardized conditions. The integrated framework allows direct comparison of assemblers presented in the study, while controlling for preprocessing and post-processing steps. Our case studies highlight the variability in assembler performance. For CLR reads, assemblers like Flye and Canu consistently produced complete bacterial genomes. For HiFi reads, Flye, Pecat, and Hifiasm yielded highly contiguous assemblies with excellent BUSCO completeness. ONT datasets with potential contamination benefited from LORA’s taxonomic analyses, enabling identification and separation of mixed species. In diploid eukaryotes, assemblers such as Necat and Pecat were able to capture haplotypic variation, albeit with increased fragmentation. Rather than ranking assemblers globally, LORA empowers users to choose based on data quality, genome complexity, and computational constraints.

### 5.4 Computational Performance and Resource Management

We quantified the computational demands of each assembler and key workflow steps (Figures 16, 17). Assembly tasks consistently dominated CPU usage, while circularization and PacBio CCS generation also contributed substantially in bacterial analyses (one third of the run time each). Memory usage varied across assemblers, with Unicycler being the most demanding, and Pecat and Necat showing the best efficiency. Parallel execution through Snakemake substantially reduces wall-clock time (i.e., real time). For instance, a complete Flye-based assembly of a streptococcus genome, including CCS generation and QC, required approximately 2 CPU hours but less than 1 hour wall-clock time when parallelized. The *Bacteroides fragilis* case study illustrates that LORA can efficiently handle both single-sample and multi-sample projects (tested with up to 50 samples), enabling large-scale comparative analyses.

**Figure 16:**
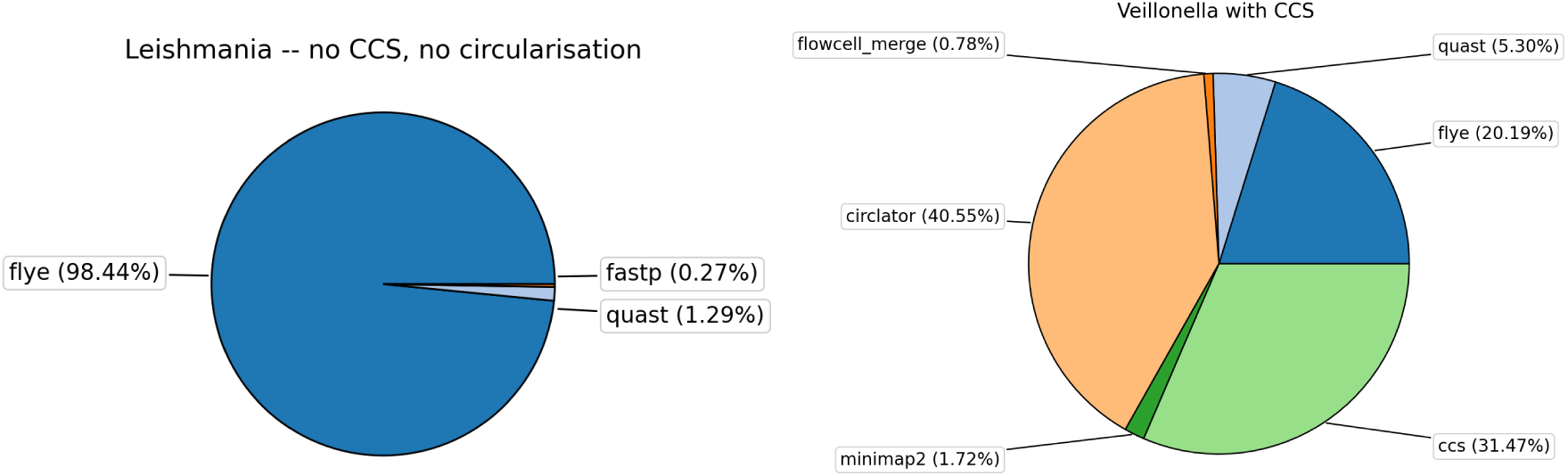
Left pie chart: distribution of CPU time for *Leishmania* data analysis using Flye assembler. Right pie chart: distribution of CPU time for the *Veillonella* data analysis from raw PacBio reads, including CCS construction and the circularization step. Tasks requiring less than two minutes of computation are not shown. When circularization and CCS preprocessing are excluded, the assembly step accounts for the majority of the runtime.

**Figure 17:**
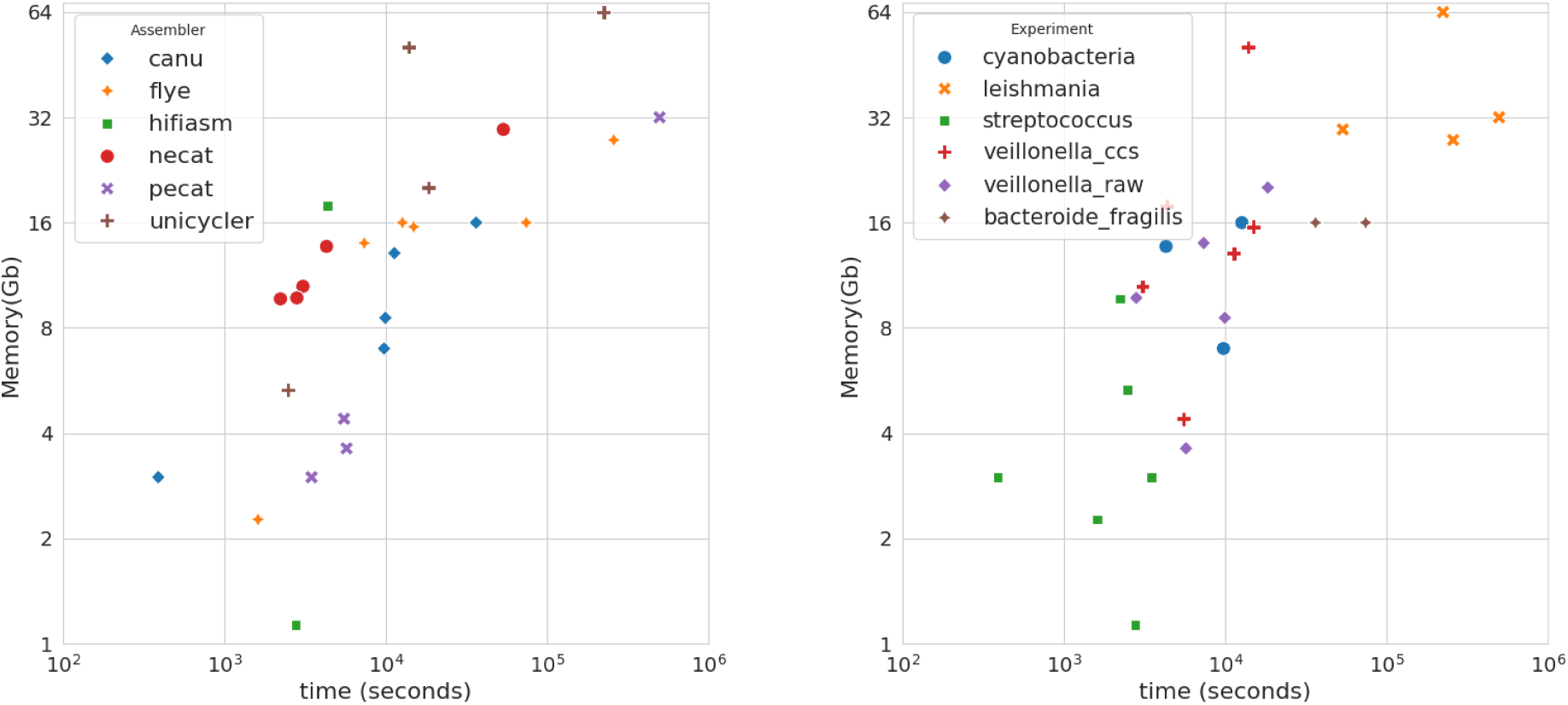
Top panel: CPU time and peak memory usage for all assembler tasks across the different experiments. Bottom panel: CPU time and peak memory usage for all experiment tasks across the different assemblers. In the bottom panel (top right corner), the *Leishmania* assemblies cluster together regardless of the assembler used, whereas on the opposite end of the spectrum lies the SGM case (HiFi data with lower yield). In the top panel, Unicycler shows substantially higher time and memory consumption, while PECAT and NECAT generally exhibit more efficient performance.

When selecting an assembler, users should balance accuracy and computational efficiency. For small bacterial genomes or constrained environments, PECAT or NECAT provide good completeness with low memory and runtime. Assemblers like Flye or Canu are more resource-intensive but yield higher contiguity and completeness, particularly for complex bacterial genomes and small eukaryotes. Unicycler may be useful for circular genomes (e.g. missing plasmid), while Hifiasm is ideal for HiFi reads and haplotype-aware assemblies.

### 5.5 Extensibility and Modularity

A key advantage of LORA lies in its modular design. The pipeline currently supports six assemblers and Illumina-based polishing with Polypolish or ONT correction (medaka). Thanks to the Snakemake architecture and containerized modules, new assemblers, polishing tools, or quality control steps can be easily integrated, supporting continuous evolution and community contributions.

### 5.6 Usability

LORA inherits its user interface from the Sequana Pipetools (https://github.com/sequana/sequana_pipetools) framework, which provides a uniform and intuitive command-line experience across all Sequana pipelines. This design ensures that common actions—such as launching the pipeline on a local machine or high-performance computing (HPC) cluster, accessing intermediate results, or resuming interrupted runs—are performed using standardized commands. Users familiar with other Sequana pipelines can therefore adopt LORA seamlessly without additional learning overhead.

In addition to these shared features, LORA introduces specific options to streamline assembly analyses. For example, the –mode option automatically configures workflow parameters for bacterial or eukaryotic genomes. When –mode bacteria is selected, the pipeline enables BUSCO and CheckM quality assessments, adjusts coverage analysis accordingly, and triggers Prokka for genome annotation. Similarly, the –data-type option standardizes parameters for different sequencing technologies—such as Oxford Nanopore or PacBio—by harmonizing assembler-specific arguments that are otherwise inconsistent across tools.

LORA also automates several tasks to simplify setup. When BUSCO or CheckM analyses are requested, the corresponding reference databases are downloaded on the fly prior to execution, ensuring analyses run smoothly without manual preparation. Before launching, LORA performs input validation to detect typographical errors and suggests corrections using a Levenshtein-distance-based algorithm. Lists of valid values and detailed option descriptions are available on the project’s wiki (https://github.com/sequana/lora/wiki ).

Together, these features make LORA accessible to users with varying levels of bioinformatics expertise, reducing configuration errors and ensuring consistent execution across computational environments.

### 5.7 Reporting

LORA also provides a comprehensive HTML report summarizing key metrics for each sample. The reports include coverage plots, BUSCO and CheckM completeness, assembly statistics, and taxonomic identifications derived from BLAST results. Although the size of HTML files may grow with large multi-sample projects, this limitation can be mitigated by generating per-sample reports or summary dashboards in future releases. Interactive assembly graphs generated by Flye can also be visualized using Bandage[45], facilitating the exploration of repeat regions and structural complexity.

## 6 Conclusion

LORA provides a flexible and reproducible framework for long-read genome assembly, integrating multiple state-of-the-art assemblers and supporting diverse sequencing technologies, including PacBio and Oxford Nanopore. Its modular design allows users to tailor workflows for bacterial or eukaryotic genomes, apply circularization when appropriate, and incorporate short-read polishing to improve assembly accuracy. By supporting haplotype-aware assemblers such as PECAT and Hifiasm, LORA can reconstruct complex diploid genomes while efficiently handling smaller bacterial replicons.

The pipeline’s polymorphic nature accommodates both single- and multi-sample experiments, while its containerized implementation ensures full reproducibility across computational environments. Through comprehensive quality assessments—including BUSCO, CheckM, N50, and coverage metrics—users can reliably evaluate assembly completeness, contiguity, and potential contamination. LORA has already been successfully applied in published studies, including Streptococcus and Veillonella genomes [6, 30], demonstrating its utility for real-world genomics projects.

By combining performance, versatility, and transparency, LORA streamlines the assembly process and provides a robust platform for benchmarking, comparative evaluation of assemblers, and reproducible genomic analyzes. Its adherence to FAIR principles and open-source design facilitates reuse, extension, and adoption by the broader research community.

Thanks to its Snakemake-based architecture and containerized modules, LORA remains easily extensible, enabling seamless integration of new assemblers and quality control tools as the field evolves.

## 7 Data Availability

All data are available in the main text or the supplementary materials.

All assemblies were performed with LORA v1.0.0, Sequana v0.19.2, Sequana Pipetools v1.3.0 and all third party tools are available on Zenodo under the Damona community as containers.

All figures and tables were generated with the Python notebooks available https://github.com/cokelaer/paper_LORA and can be executed online (DOI 10.5281/zenodo.17250176)

## 8 Acknowledgments

The authors thank Muriel Gugger (Institut Pasteur, Paris) for sharing the unpublished cyanobacterial dataset used in this study; Christophe Beloin (Institut Pasteur, Paris) for authorizing the use of the Veillonella BAM dataset; and Shaynoor Dramsi (Institut Cochin) for authorizing the use of the Streptococcus BAM dataset. The authors also thank the Biomics platform (Institut Pasteur, Paris, France) for sequencing the *Veillonella*, *Streptococcus*, and Cyanobacterial data sets.

## Annex

### 8.1 Veillonella Parvula case

In this example, the PacBio dataset contains 338,310 raw reads, with a mean read length of 9 kb and an N50 of 13 kb. As described in the main text, we also generated corrected reads (CCS). Since this is an older dataset, CCS reads were produced from the raw data using non-stringent parameters (a low quality threshold of RQ=0.7 and no restriction on the number of passes). This process yielded 192,532 CCS reads, and after filtering out reads shorter than 1 kb, 174,520 reads remained for assembly. Figure 18 shows the distribution of corrected reads as a density plot of read length versus GC content. The mean length is 8.8 kb and the mean GC content is approximately 40%, as expected for this species. This genome belongs to class I, as illustrated in Figure 19, where the maximum repeat length is below 2 kb.

**Figure 18:**
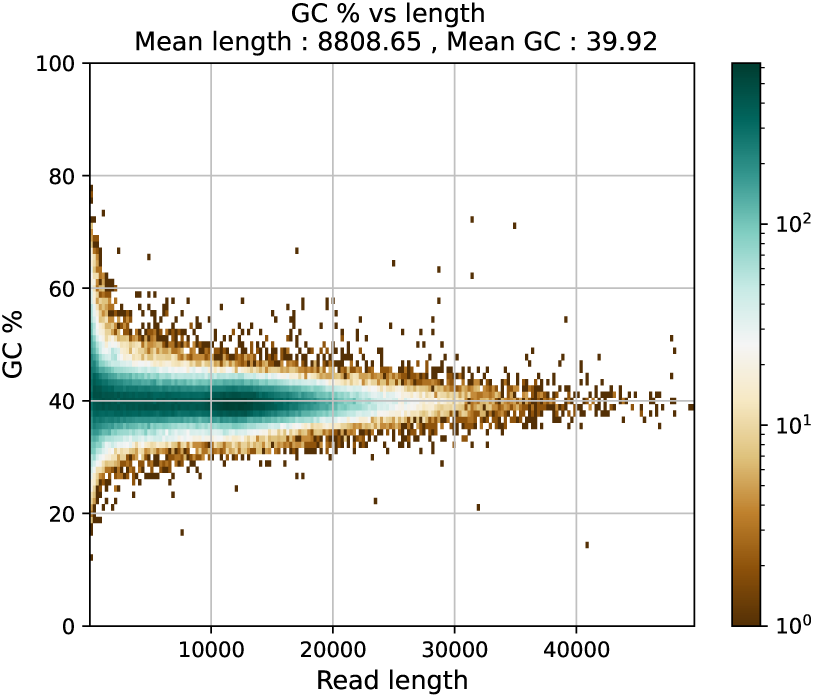
2-dimensional density of read length versus GC content. dark blue dots indicates higher densities. *Veillonella parvula*) data set.

**Figure 19:**
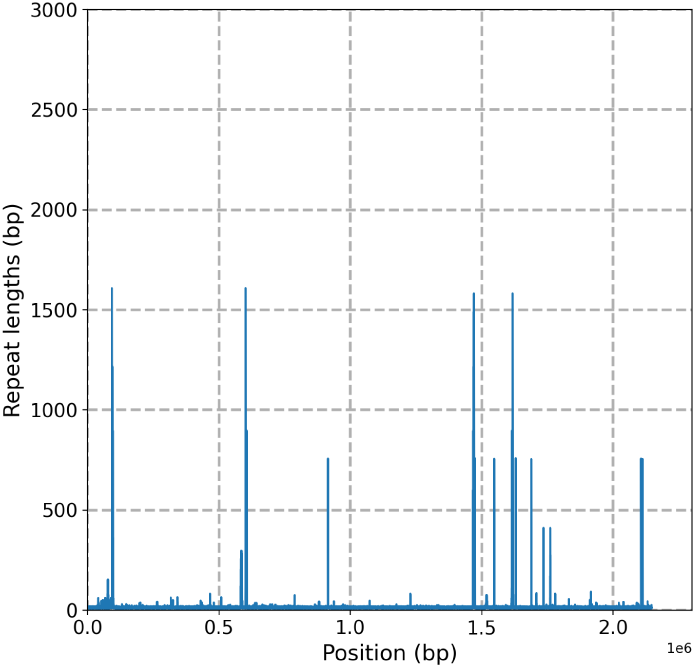
Repeats length and position along the genome of Veillonella Parvula. Class I genome with few repeats below 2kb.

To analyze this dataset, we used the LORA command shown in Listing 1. The corresponding workflow is presented in Figure 20.

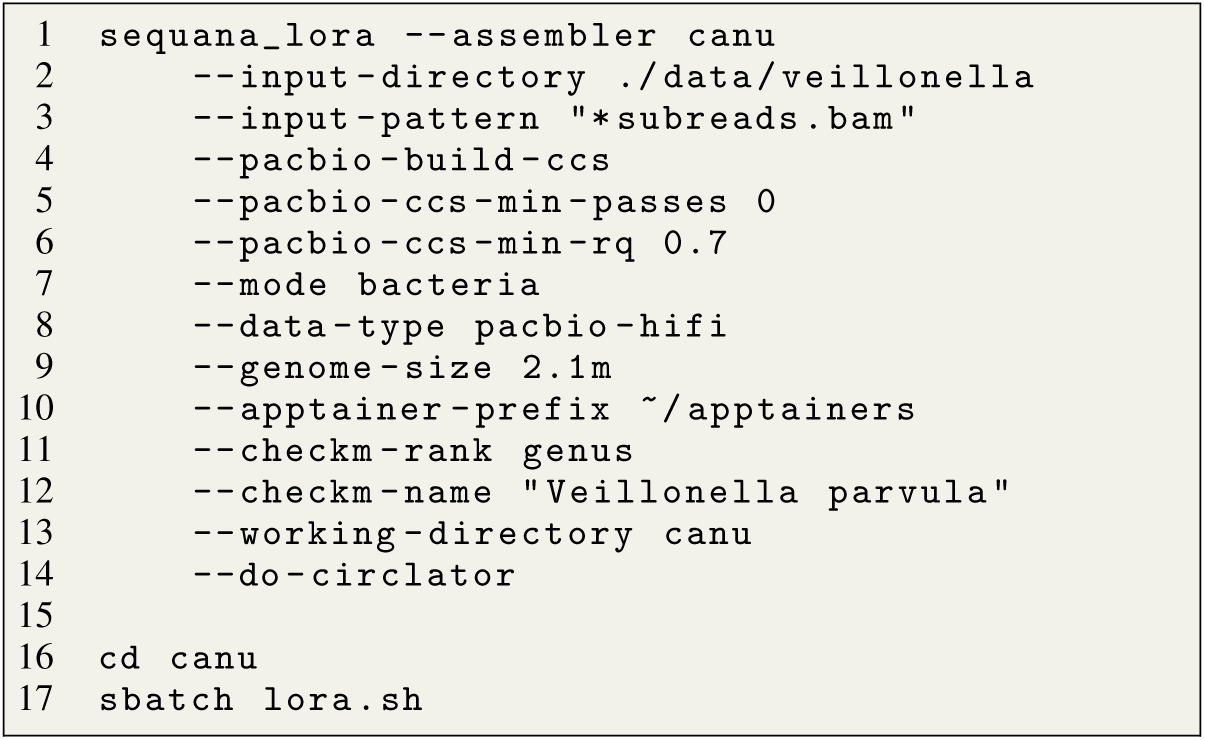

Listing 1: This script analyzes the Veillonella dataset, generating CCS reads (line 4) and assessing genome completeness with CheckM (lines 10–122).

**Figure 20:**
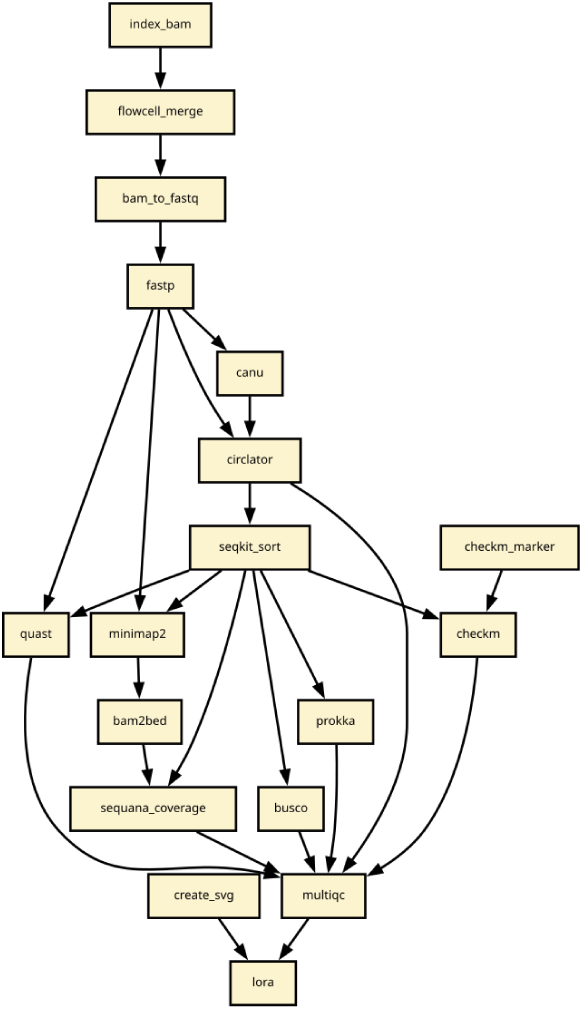
Workflow used in the Veillonella Parvula case following command in Listing 1

### 8.2 Streptococcus gallolyticus subsp. macedonicus

In Figure 21, we show the distribution of the CCS reads of the *Streptococcus gallolyticus* data set, illustrating the read length versus GC content. We built the CCS data setting HiFi parameters (10 passes, accuracy of 0.99). The mean length is 7.3 kb and GC is about 38% as expected for that species. HiFi data is made of 34,013 reads. Using the same parameters as in [30], with CCS using 3 passes retained, we would obtain 153,120 reads of 11.7 kb read length. The completeness (CheckM) could be as large as 99.96%. The analysis of the repeats within the assembled genome reveals a large 10 kb repeat and several 5 kb repeats. This genome belongs to the class III bacterial genome. An example of the pipeline used for this data set (HiFi, CCS build, circularisation) is shown in Figure 23 and the code used to prepare the LORA analysis working directory is provided in Listing 2.

**Figure 21:**
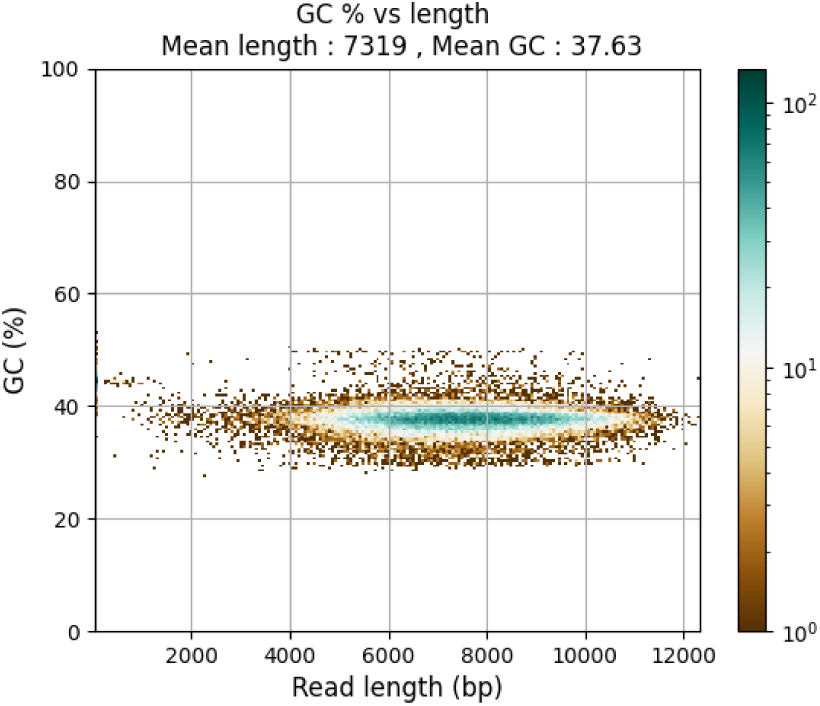
2-dimensional density of read length versus GC content. dark blue dots indicates higher densities. *SGM*) .

**Figure 22:**
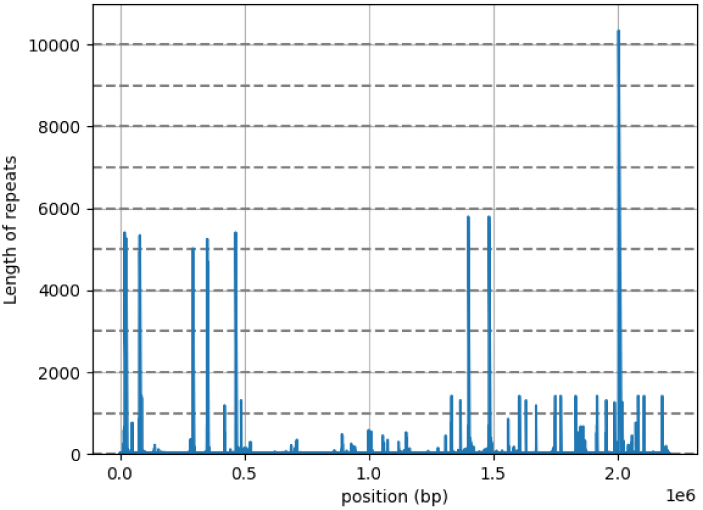
Repeats length and position along the genome of *Streptococcus gallolyticus*. Class III genome with presence of repeats *≥* 7*kb*

**Figure 23:**
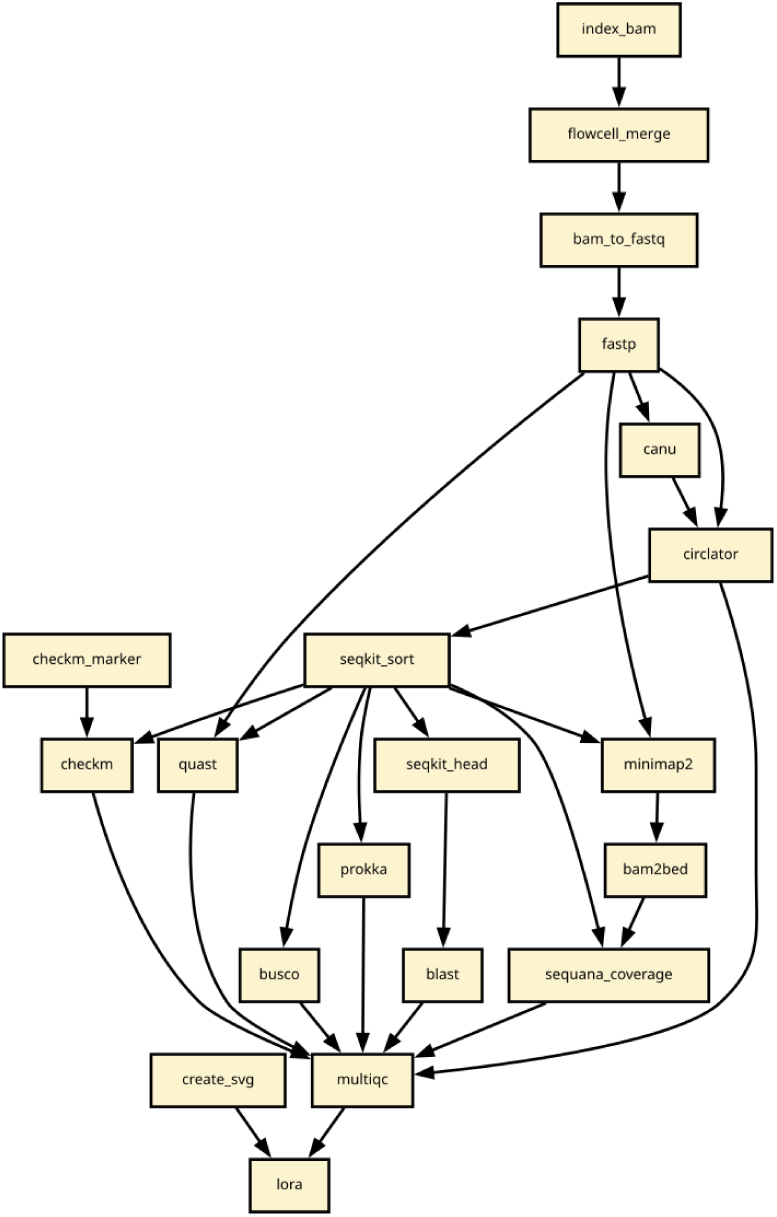
Workflow used in the SGM analysis (assembler set to Canu, and circularisation on). Since input data is in BAM format, extra rules are added in the top of the workflow.

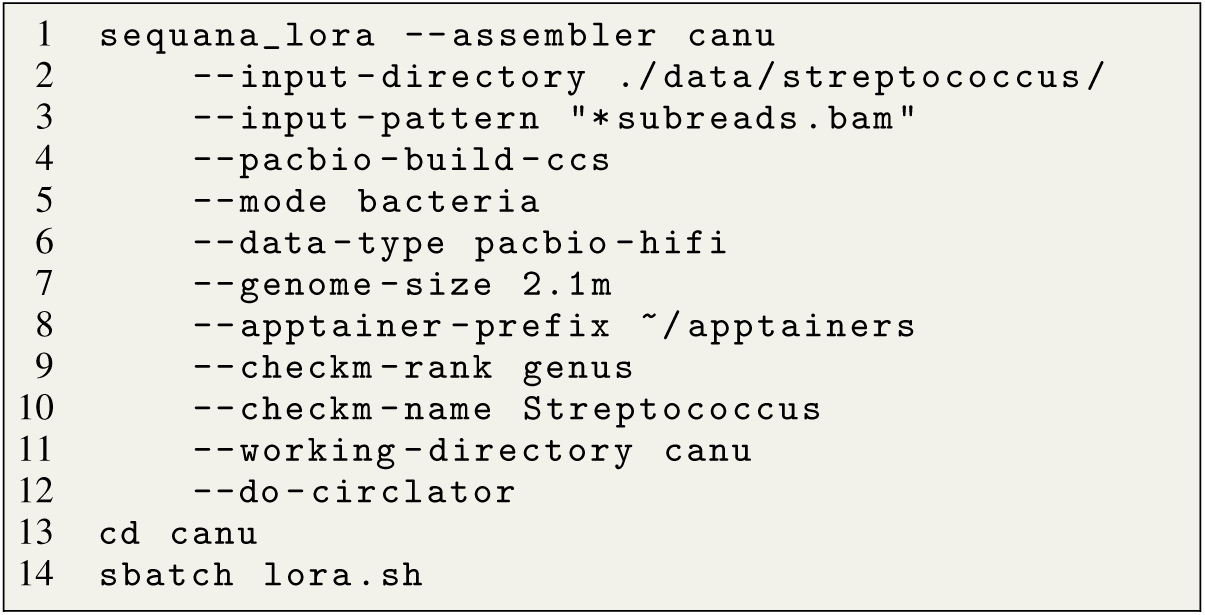

Listing 2: script used to analyse *Streptococcus* data. on line 4, we build the CCS data. One line 6, we specify the type of data. One line 12, we can add an optional circularisation.

### 8.3 Cyanobacterium

Based on genome reference assembly obtained in this paper, we can look at the repeats of the two mixed strains. We can also look at GC versus read length

**Figure 24:**
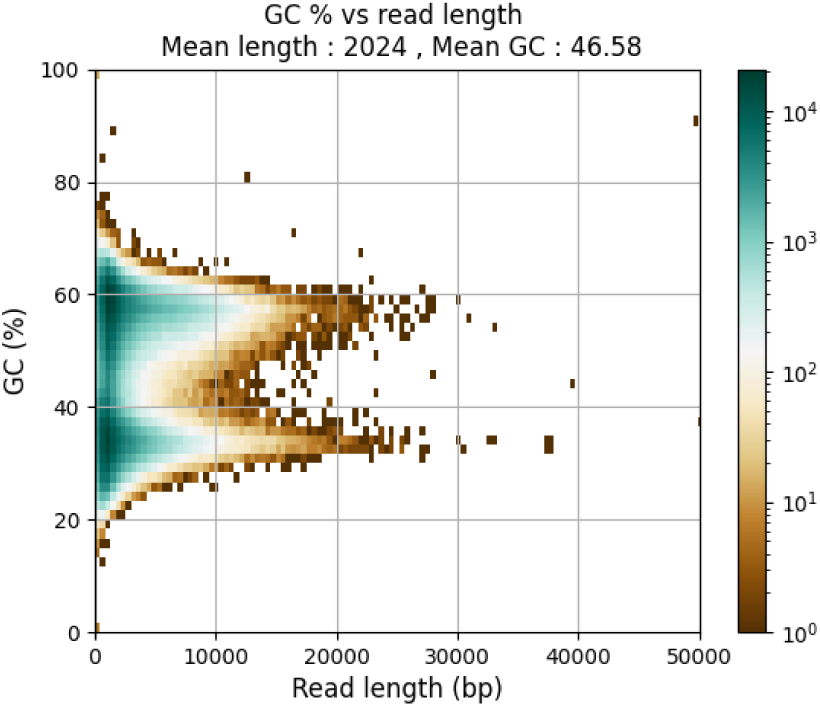
2-dimensional density of read length versus GC content. dark blue dots indicates higher densities. Nanopore data Cyanobacterium

**Figure 25:**
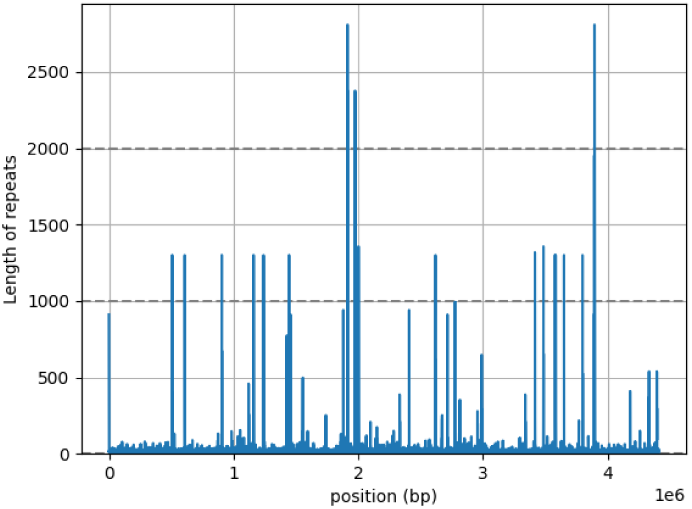
Repeats length and position of the first cyanobacterial genome (class I).

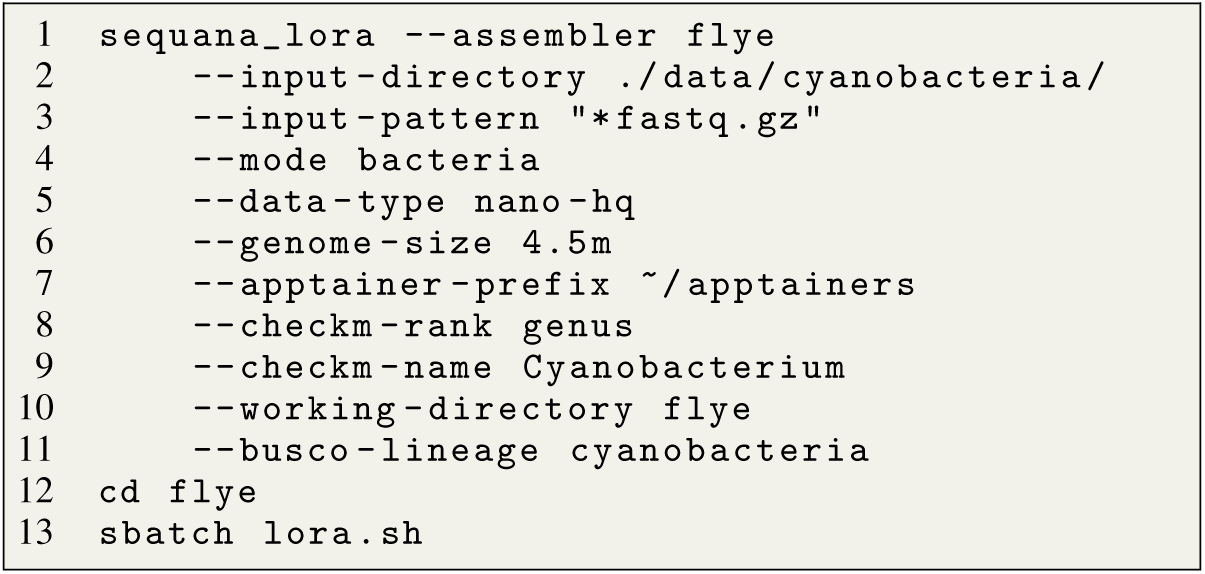

Listing 3: script used to analyse *Cyanobacterium* data. on line 4, we build the CCS data. One line 6, we specify the type of data. One line 12, we can add an optional circularisation.

**Figure 26:**
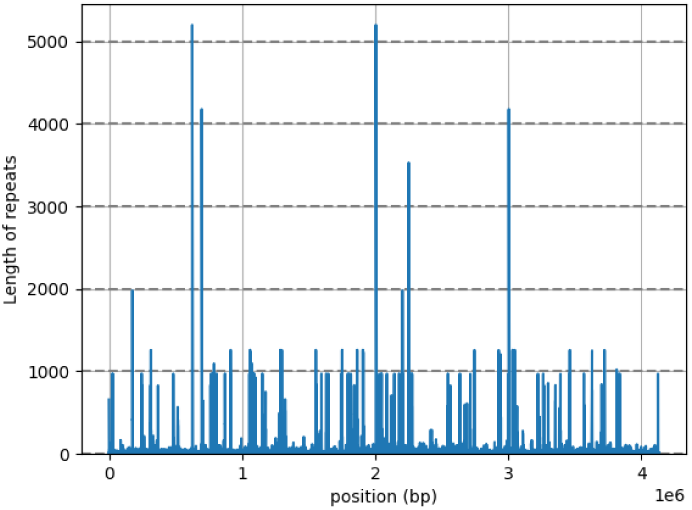
Repeats length and position of the second cyanobacteria (class I)

**Figure 27:**
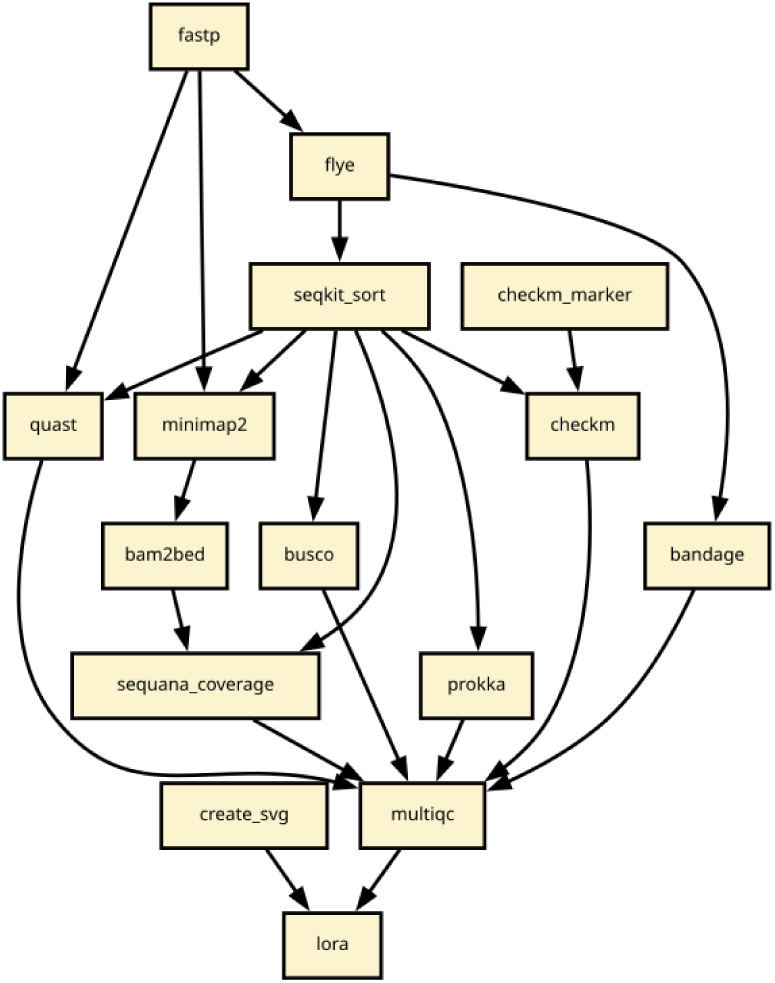
Workflow used in the Cyanobacteria analysis (assembler set to Canu, and with no circularisation on. because the assembler is set to flye, the bandage rule was added automatically (visualisation of flye underlying assembly graph.).

### 8.4 Bacteroides Fragilis

Based on genome reference GCF 000025985.1 (ASM2598v1), we computed the repeats shown in Figure 29 and present the distribution of the raw Nanopore reads. The mean length is 5.97 kb and the GC content is about 44% as expected for that species.

**Figure 28:**
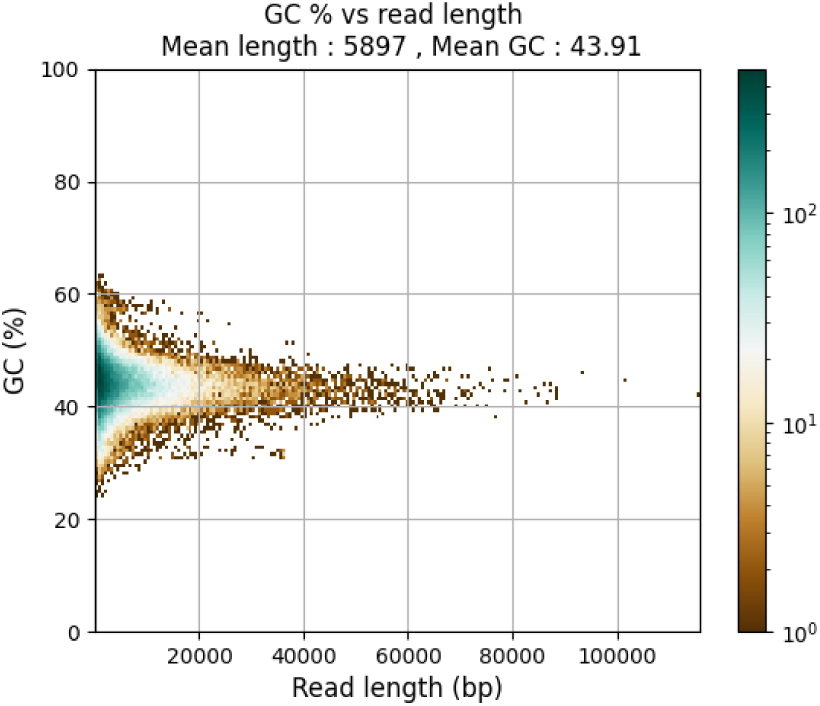
2-dimensional density of read length versus GC content. dark blue dots indicates higher densities. Nanopore data sample CGU5846T.

**Figure 29:**
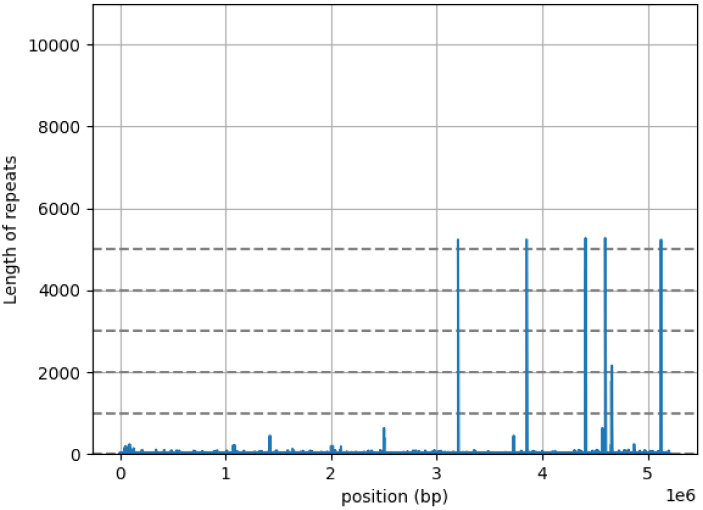
Repeats length and position along the genome of *Bacteroides Fragilis*. Class I genome with few repeats in 5-7 kb.

**Figure 30:**
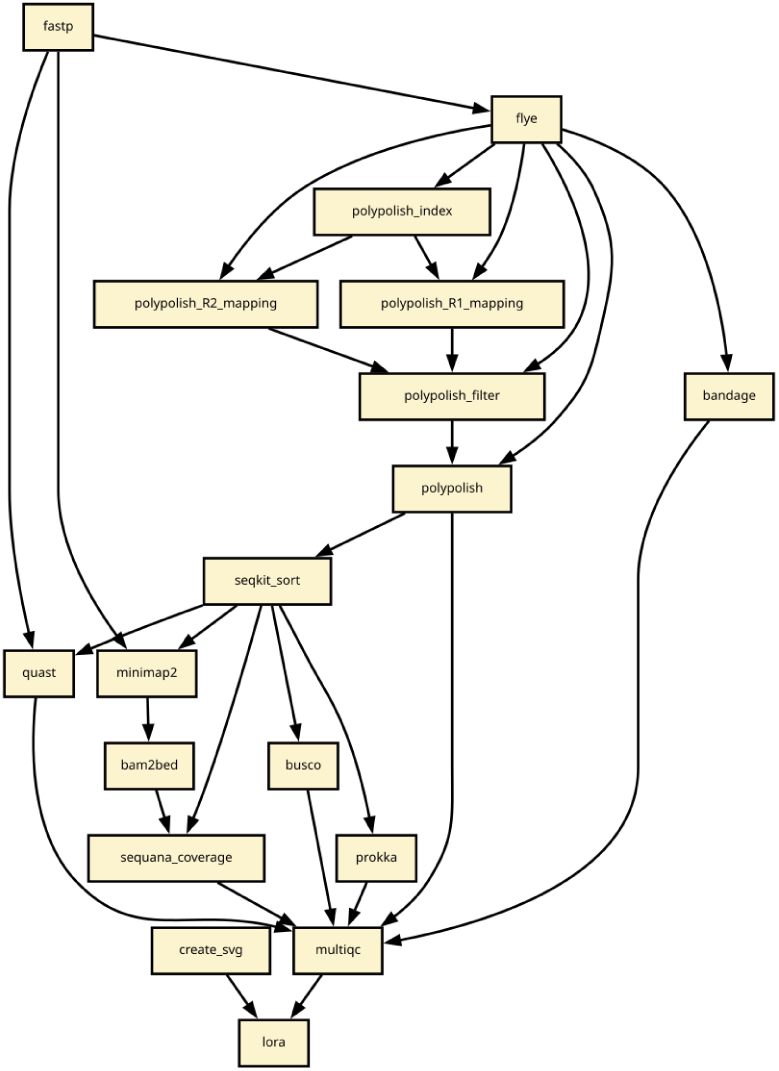
Workflow used in the the *Bacteroides fragilis* case. Polishing was added, which adds 5 rules.

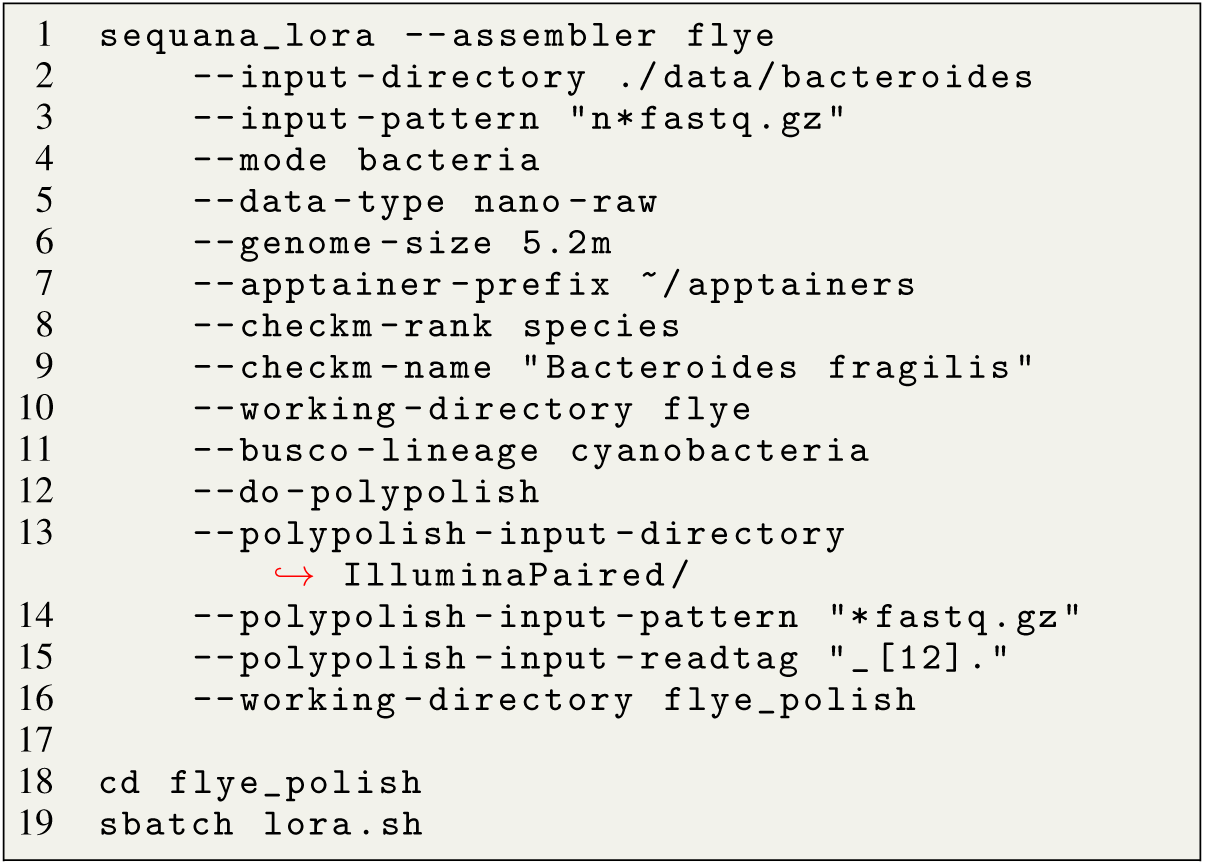

Listing 4: script used to analyse *Bacteroides fragilis* data. Given Illumina data, we can polish the data (line 12-15)

### 8.5 Leishmania case

**Figure 31:**
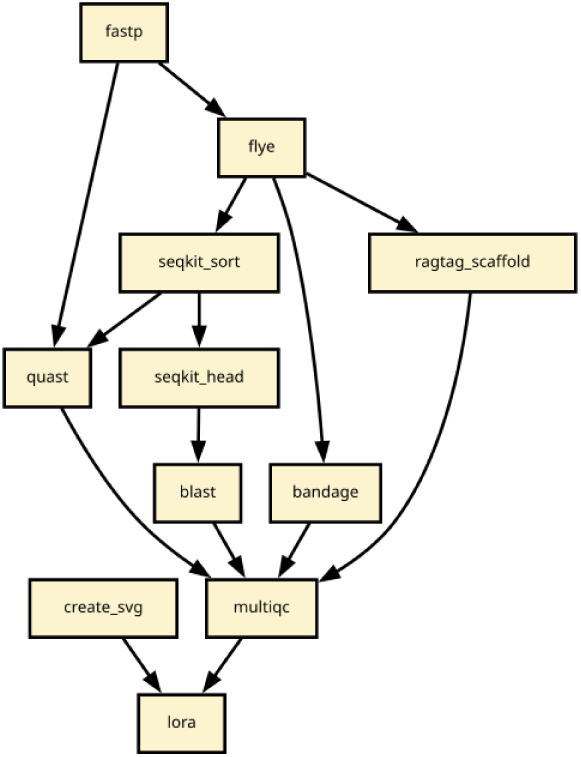
Workflow used in the *Leishmania* case

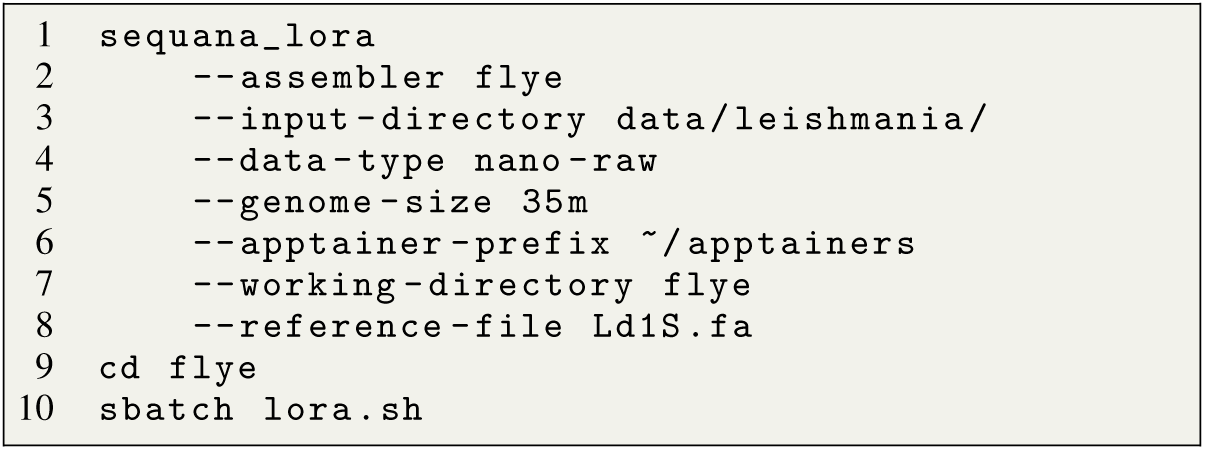

Listing 5: script used to analyse Leishmania data.

